# The diversity of *SNCA* transcripts in neurons, and its impact on antisense oligonucleotide therapeutics

**DOI:** 10.1101/2024.05.30.596437

**Authors:** James R. Evans, Emil K. Gustavsson, Ivan Doykov, David Murphy, Gurvir S. Virdi, Joanne Lachica, Alexander Röntgen, Mhd Hussein Murtada, Chun Wei Pang, Hannah Macpherson, Anna I. Wernick, Christina E. Toomey, Dilan Athauda, Minee L. Choi, John Hardy, Nicholas W. Wood, Michele Vendruscolo, Kevin Mills, Wendy Heywood, Mina Ryten, Sonia Gandhi

**Affiliations:** Department of Clinical and Movement Neurosciences, UCL Queen Square Institute of Neurology, University College London; London, UK; The Francis Crick Institute; London, UK; Aligning Science Across Parkinson’s (ASAP) Collaborative Research Network; Chevy Chase, MD, 20815; Genetics and Genomic Medicine, Great Ormond Street Institute of Child Health, University College London; London, UK; UK Dementia Research Institute at The University of Cambridge, Cambridge, United Kingdom; Queen Square Brain Bank for Neurological Disorders, UCL Queen Square Institute of Neurology; London, UK; Yusuf Hamied Department of Chemistry, University of Cambridge; Cambridge, UK; Department of Brain & Cognitive Sciences, KAIST, 921 Dehak-ro, Daejeon, Republic of Korea; Department of Neurogenerative Disease, UCL Queen Square Institute of Neurology; London, UK; UCL Dementia Research Institute, University College London, London, UK; NIHR Great Ormond Street Hospital Biomedical Research Centre, University College London; London, UK; Department of Clinical Neurosciences, School of Clinical Medicine, The University of Cambridge, Cambridge, UK

**Keywords:** *SNCA*, antisense oligonucleotide, neuron, Parkinson’s disease, long-read, transcripts, αSyn

## Abstract

The role of the *SNCA* gene locus in driving Parkinson’s disease (PD) through rare and common genetic variation is well-recognized, but the transcriptional diversity of *SNCA* in vulnerable cell types remains unclear. We performed *SNCA* long-read RNA sequencing in human dopaminergic neurons and show that annotated *SNCA* transcripts account for only 5% of expression. Rather, the majority of expression (75%) at the *SNCA* locus originates from transcripts with alternative 5’ and 3’ untranslated regions. Importantly, 10% originates from transcripts encoding open reading frames not previously annotated, which are translated and detectable in human postmortem brain. Defining the 3’ untranslated regions enabled the rational design of antisense oligonucleotides targeting the majority of *SNCA* transcripts, leading to the effective reversal of PD pathology, including protein aggregation, mitochondrial dysfunction, and toxicity. Resolving the complexity of the *SNCA* transcriptional landscape impacts RNA therapies and highlights differences in protein isoforms and their contribution to disease.

## Introduction

Parkinson’s disease (PD) is an incurable neurodegenerative disease, characterized by the progressive loss of midbrain dopaminergic (mDA) neurons and the aggregation of the protein α-synuclein (αSyn; encoded by the *SNCA* gene), which typically accumulates with lipid membranes to form neuronal inclusions known as Lewy bodies.^1,2^ Substantial genetic and pathological evidence indicates that αSyn aggregation is a primary driver of neuronal death.^3,4^

Quantitative and qualitative differences in αSyn are causally linked to PD. Chronic overexpression of αSyn, due to multiplications of the *SNCA* locus, is a well-defined cause of parkinsonism and dementia.^5,6^ Patients with triplications of the *SNCA* locus develop early-onset disease with rapidly progressive symptoms, whereas patients with duplications develop late-onset parkinsonism, and may also remain non-penetrant in advanced age,^6–8^ suggesting a gene dosage effect. Additionally, several single-nucleotide polymorphisms (SNPs) at the *SNCA* locus are associated with both increased susceptibility to idiopathic PD (iPD) and increased *SNCA* expression.^9,10^ *SNCA* missense mutations also cause familial PD and alter the aggregation kinetics and lipid binding of αSyn. For example, *SNCA* (c.209G>A; A53T) causes late-onset autosomal dominant PD with a more aggressive course than that of iPD.^11,12^

Three shorter αSyn protein-coding transcripts (αSyn-126, αSyn-112, and αSyn-98), in addition to the full-length transcript (αSyn-140), have been reported,^13^ presenting another mechanism by which qualitative variation in αSyn could be generated. In fact, evidence suggests that each isoform is associated with differing aggregation propensity,^14–16^ differential expression in Lewy body disorders^14,17^ and variable usage in MPP^+^ and rotenone models.^18^ Furthermore, outside of the open reading frame (ORF), alternative 3’ untranslated region (UTR) usage has been demonstrated to affect mRNA stability and localisation in neurons,^19^ and an extended *SNCA* 3’UTR may play a pivotal role in regulating αSyn expression levels^20,21^ and localisation.^22^ These are likely critical features in PD given that small fluctuations in αSyn concentration, or isoform usage, may alter its propensity to aggregate.^4,16^ In this context, it is surprising that the diversity in *SNCA* transcription has not received more attention, particularly given that a recent study in post-mortem brain demonstrated far more transcript complexity at the locus than previously appreciated.^23^

Antisense oligonucleotides (ASOs) have recently emerged as a promising therapeutic strategy across neurodegenerative disease^24^ due to their capability to affect both the quantity and quality of RNA. In the context of PD, ASOs aiming to reduce total αSyn protein offer key advantages over similar αSyn-targeting strategies, as they target intracellular pools of *SNCA* mRNA and ultimately protein, and therefore intracellular aggregation. ASOs targeting *SNCA* quantitatively have already shown promise in animal models,^25–30^ and are currently in clinical trial for a related synucleinopathy multiple system atrophy (NCT04165486).

Without a detailed understanding of the landscape of *SNCA* transcripts in human dopaminergic neurons, RNA targeting approaches may prove to be less effective. We perform targeted long-read RNA-seq in control and patient enriched iPSC-derived mDA neurons to uncover the transcriptional diversity of the *SNCA* gene in health and disease, and validate these transcripts in postmortem human brain tissue. Using this patient-derived neuronal model, which captures the human synucleinopathy *in vitro,*^31^ we test a *SNCA*-targeting ASO approach and demonstrate effective reduction in *SNCA* transcript expression and protein aggregation, and a reversal in established PD-associated cellular pathology.

## Results

### Novel *SNCA* transcript structures are detected in iPSC-derived midbrain dopaminergic neurons

The diversity of *SNCA* transcripts in iPSC-derived mDA neurons is currently not known. We generated mDA neurons from iPSCs using an optimised small molecule protocol (**Fig. 1A**),^31^ which generates highly enriched mDA neurons, with 91% cells expressing the neuronal marker MAP2, and 87% cells expressing the mDA neuronal marker TH by ICC (**Fig. 1B-C**). A significant upregulation of mDA neuronal maturation markers *TH* and *SLC6A3*, a dopamine transporter gene, occurred in mDA neuronal cultures compared to mDA-specific neural precursor cells (NPCs) by qPCR (**Fig. 1D**).

**Fig. 1.**
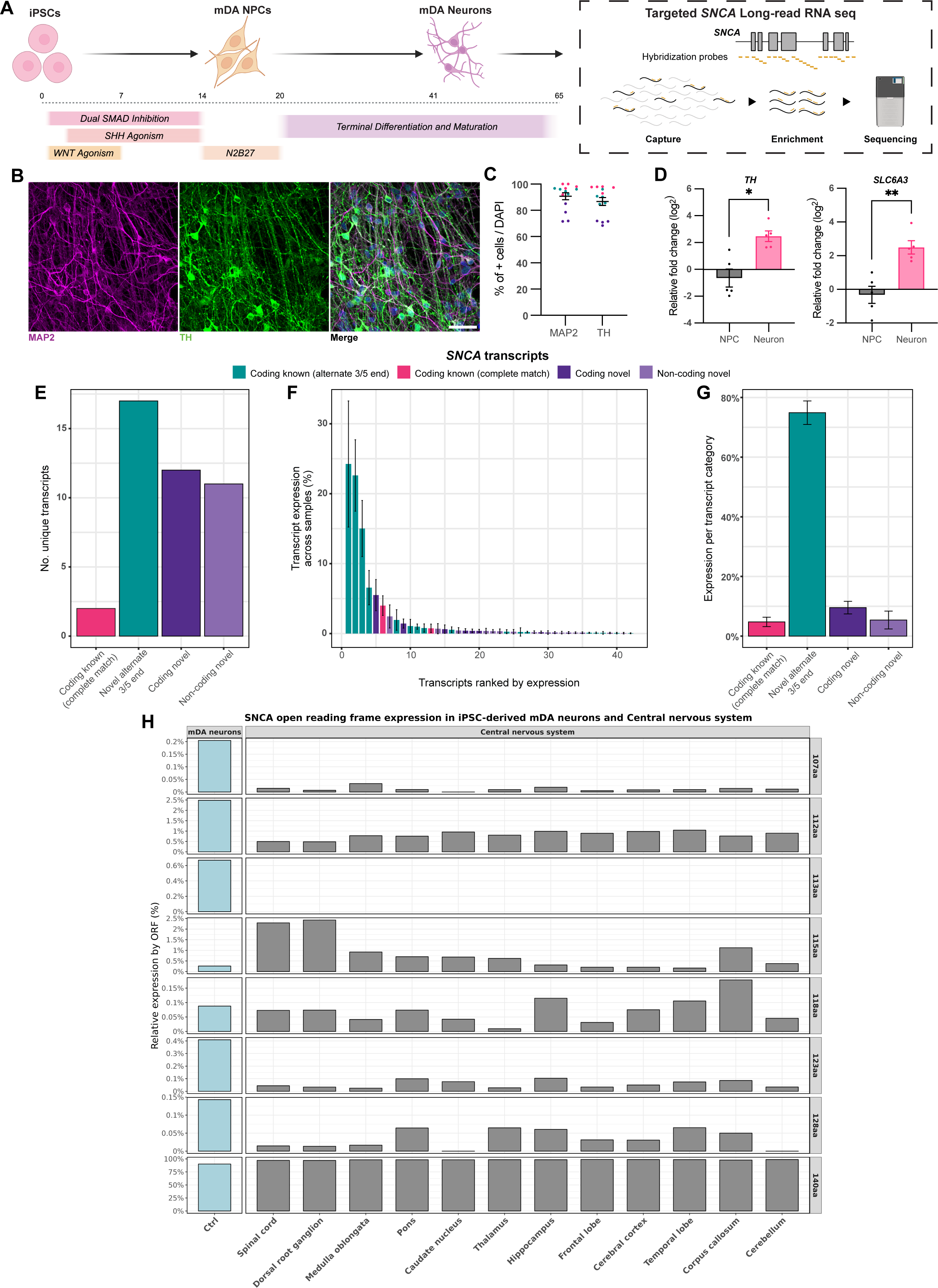
*SNCA* transcripts in enriched control iPSC-derived mDA neurons. (**A**) Schematic outlining mDA differentiation protocol and targeted long-read RNA-seq paradigm. (**B**) Representative ICC images showing MAP2 and TH expression after 65 days of differentiation (scale bar = 40 μm). (**C**) ICC quantification showing that 91% of cells express MAP2 and 87% of cells express TH. (**D**) Quantitative PCR showing mRNA expression of *TH* and *SLC6A3* in mDA neurons relative to mDA NPCs (\**P* < 0.05, paired t-test). (**E**) Bar chart depicting number of unique *SNCA* transcripts identified per transcript category through targeted long-read RNA-seq in control iPSC-derived mDA neurons. (**F**) Normalized expression per *SNCA* transcripts corresponding to the percentage of expression per transcripts out of total expression of the loci. (**G**) Bar chart depicting the percent expression of *SNCA* per transcript category. (**H**) Relative open reading frame expression in control iPSC-derived mDA neurons and across postmortem brain regions.

Using enriched neuronal cultures from control iPSCs (3 donors), we performed targeted Pacific Biosciences (PacBio) long-read isoform sequencing (Iso-Seq). cDNAs were enriched using 111 biotinylated hybridization probes designed against exonic (n = 41) and intronic (n = 70) genic regions of *SNCA* according to annotation (GENCODEv38) (**Fig. S1**) to generate a mean of 1,197 ± 939 (range = 74-3,485) full-length HiFi reads per sample. Collapsing mapped reads resulted in the identification of 283 unique *SNCA* transcripts after quality control. We filtered transcripts based on their use (≥0.1% transcript usage per sample) and identified 42 unique *SNCA* transcripts (**Fig. 1E, Fig. S2**), each supported by a mean of 863 ± 2,203 (range = 24-10,936) full-length HiFi reads per transcript, providing a detailed annotation of *SNCA* transcription in an enriched single-cell type that is critical to PD pathology.

Of these unique transcripts only 2 precisely matched transcripts already in annotation (Coding known (complete match)). We identified a further 17 transcripts with alternative 5’ and 3’UTRs but with an ORF matching annotation (Novel alternate 3’/5’ UTR) including 3 matching αSyn-112 (CCDS43252) and 14 matching αSyn-140 (CCDS3634). Finally, we identified 12 transcripts with novel ORFs (Novel coding) and 11 novel non-coding transcripts (Novel non-coding). It is notable that we did not identify any transcripts matching the commonly studied αSyn-126 and αSyn-98 in this model.

After ranking transcripts by their level of relative expression we found that the most highly used transcript accounted for only 24.3 ± 9.0%, demonstrating that there was no single dominant transcript in mDA neurons (**Fig. 1F**). Interestingly, the two transcripts completely matching annotation, represented only 5% of *SNCA* expression. Most transcription at the *SNCA* locus originated from 17 (75%) transcripts with known open reading frames (ORFs) but novel 5’ and 3’ untranslated regions (UTRs). It was noteworthy that the 12 transcripts with novel ORFs accounted for 10% of expression (**Fig. 1G**). Overall, we identified seven novel ORFs (αSyn-107, αSyn-112, αSyn-113, αSyn-115, αSyn-118, αSyn-123, and αSyn-128), corresponding to these 12 transcripts. We sought to validate the novel coding transcripts in human brain. Using long-read RNA-seq we detected all (αSyn-107, αSyn-112, αSyn-115, αSyn-118, αSyn-123, and αSyn-128) but one (αSyn-113) of the novel ORFs across 12 brain regions derived from an average of 25 ± 9 (range 3-47) donors per brain region (**Fig. 1H**).

### Novel protein coding *SNCA* transcripts are translated in brain

Given the detection of these novel ORFs in human postmortem brain and the fact that αSyn protein quantity and quality influences its aggregation propensity, we focused further on protein coding isoforms. These novel ORFs differed from the most commonly studied and abundant αSyn isoform (αSyn-140) due to partial loss of the first coding exon (αSyn-118, αSyn-123 and αSyn-128), an extension of the second coding exon (αSyn-118 and αSyn-128), an extension (αSyn-115) or a partial loss (αSyn-107 and αSyn-113) of the third coding exon, a loss of the fourth coding exon (αSyn-107, αSyn-112 and αSyn-115) or a loss of the last coding exon (αSyn-107 and αSyn-115). For one isoform (αSyn-107) there is an extension of the ORF to include part of the 3’UTR (**Fig. 2A**). Three ORFs (αSyn-107, αSyn-118 and αSyn-128) were predicted to contain amino acid sequences that are absent from annotation (both currently annotated isoforms of *SNCA* or within the UniProt database).

**Fig. 2.**
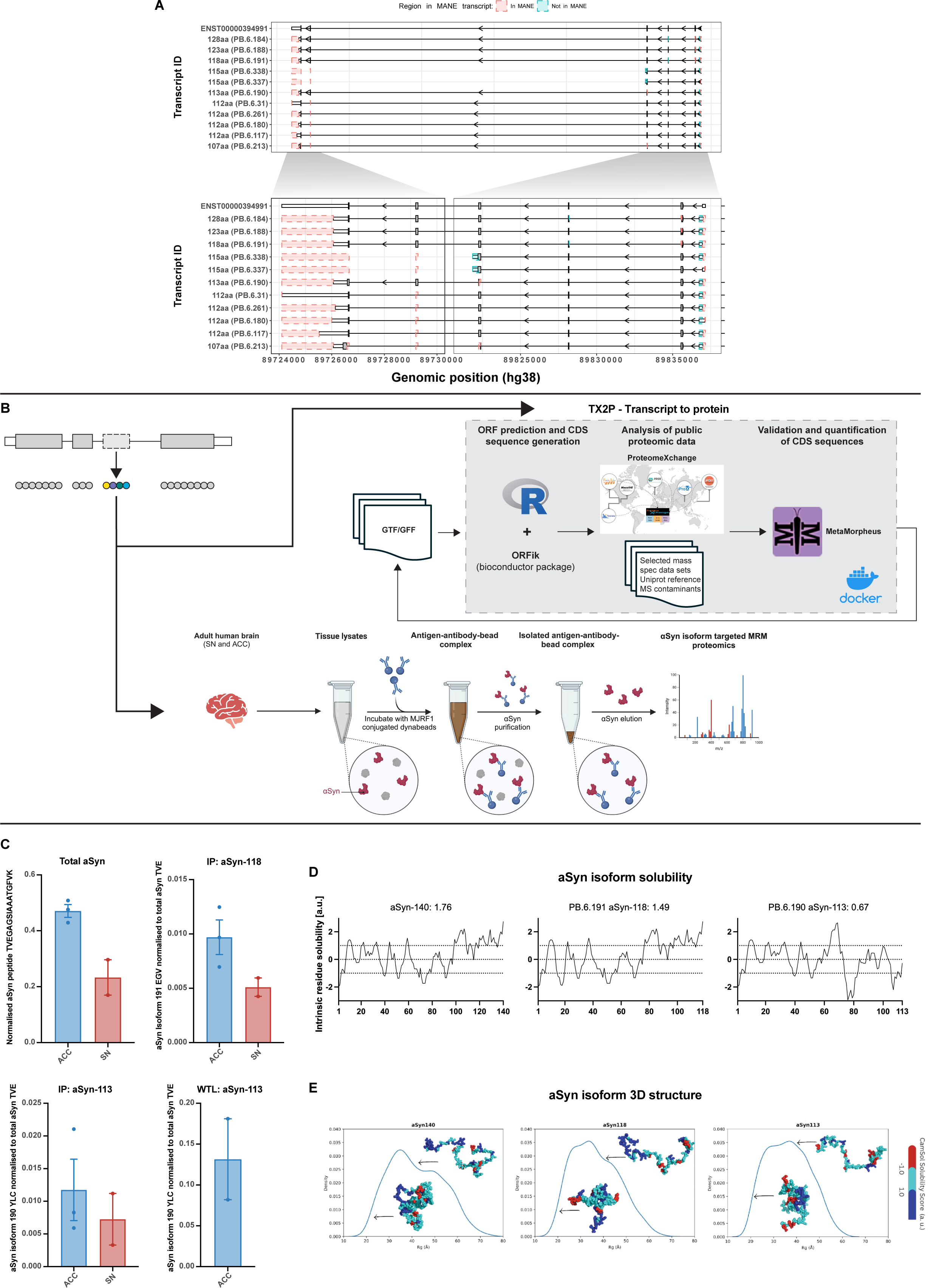
Novel coding *SNCA* transcripts are translated. (**A**) Novel coding *SNCA* transcripts plotted using ggtranscript with differences as compared to MANE select (ENST00000394991) highlighted in blue and red. (**B**) Schematic of the strategy used to detect translation of novel coding *SNCA* transcripts through αSyn isoform targeted MRM proteomics and TX2P. (**C**) Abundance in tissue immunoprecipitation of total αSyn, abundance of αSyn-118 ratioed to total αSyn, and αSyn-113 detected in immunoprecipitation and whole tissue lysate ratioed to total αSyn. (**D**) Solubility plots of αSyn-140, αSyn-118, and αSyn-113 using the CamSol method. (**E**) Conformational ensembles using AlphaFold of αSyn-140, αSyn-118, and αSyn-113.

Given our high confidence in these novel ORFs at the RNA level, we investigated their putative role in PD by assessing the transcript usage of the seven novel ORFs in patient iPSC mDA neurons. Using two different mutation types, a patient line with a triplication of the *SNCA* locus (SNCAx3) and a missense mutation (A53T), we found that the relative usage of αSyn-118 (PB.6.191) was significantly higher in iPSC-derived mDA neurons from SNCAx3 compared to controls (**Fig. S4B**) suggesting that the pathogenicity of the locus triplication could be mediated by both qualitative and quantitative changes in αSyn.

Detection of these novel ORFs is challenging due to the low abundance of the newly identified isoforms and reliance on the identification of unique peptide sequences in mass spectrometry-based approaches. Of the 8 novel ORFs, we noted that five generated unique peptide sequences (not present in annotated SNCA isoforms or other proteins within UniProt) (**Fig. S5A-B**). Importantly, these peptide sequences could not be generated through cleavage of αSyn protein. We investigated if these unique tryptic peptides (representing the three isoforms) were detectable using targeted proteomic multiple reaction monitoring (MRM) LC-MS/MS method in immunoprecipitated αSyn from post-mortem brain tissue from two PD brain regions from 3 individuals (**Fig. 2B**). This approach detected αSyn-118 and αSyn-113 in both brain regions in the αSyn enriched fractions, and we detected αSyn-113 in whole brain lysate (**Fig. 2C, Fig. S6-7**).

Subsequently, we investigated publicly available mass spec data (**Supplementary table 8**) using TX2P, a software tool to screen for unique peptide sequences based on long-read RNA structures. We were able to show the translation of αSyn-128 through the identification the amino acid sequence EGVVAAAEKTKQGVFVGSK in human prefrontal cortex (Broadman area 9) (Q value <0.01). This data set consists of human brain tissue collected post-mortem from patients diagnosed with multiple system atrophy (n = 45) and from controls (n = 30).^32^ Overall, these approaches provided translational support for 3 novel isoforms in human brain.

Once translated, these novel coding isoforms may exhibit different behaviour to the known ORF 140 amino acid transcript. The aggregation propensity of any isoform is dependent on several characteristics: the amino acid sequence, and biophysical properties of the residues; the exposure of aggregation promoting residues in the structure; and the thermodynamic stability of the native state. Moreover, the solubility and the aggregation rate of a protein are related, and both dependent on the biophysical properties of the amino acid sequence. We adopted the CamSol method which first calculates a solubility profile (a score for each residue in the sequence), and then calculates an overall solubility score from the profile itself.^33,34^ We found that certain isoforms, including αSyn-118, have a lower CamSol score than the canonical αSyn-140, predicting a lower solubility and thus a higher propensity to aggregate (**Fig. 2D**). We also demonstrate the predicted structures of these isoforms (**Fig. 2E**) that indicate potential hotspots of low solubility that may lead to αSyn self-assembly. Thus, in the SNCAx3 state, the increased transcript use of a more insoluble isoform of alpha-synuclein is likely to impact the aggregation and pathology observed.

### Targeting *SNCA* RNA through the rational design of *SNCA*-targeting ASOs

*SNCA* RNA targeting approaches have the potential to modify disease, in particular, ASO therapies aimed to reduce the quantity of *SNCA RNA* by triggering RNase H1-dependent RNA degradation. Effective ASO design requires consideration of target sequence, chemical modifications, and the time-course of delivery and measurement of response.

We focussed the ASO design on the most unique region of *SNCA*, namely the 3’UTR, and designed nine 20mer ASOs targeted across the 3’UTR of *SNCA* (table) with all sequences modified to include phosphorothioate (PS) linkages throughout. These nine ASOs were screened in SH-SY5Y cells by transfection of the nine *SNCA*-targeting ASOs and a control non-targeting ASO (ASO-Ctrl) and quantification of *SNCA* expression after 24- and 48-hours using qPCR. Four ASOs (ASO-1, ASO-3, ASO-7, and ASO-9) significantly reduced *SNCA* expression compared to those treated with ASO-Ctrl after 24 hours (**Fig. 3A**), but this was not maintained after 48 hours (**Fig. S8A**).

**Fig. 3.**
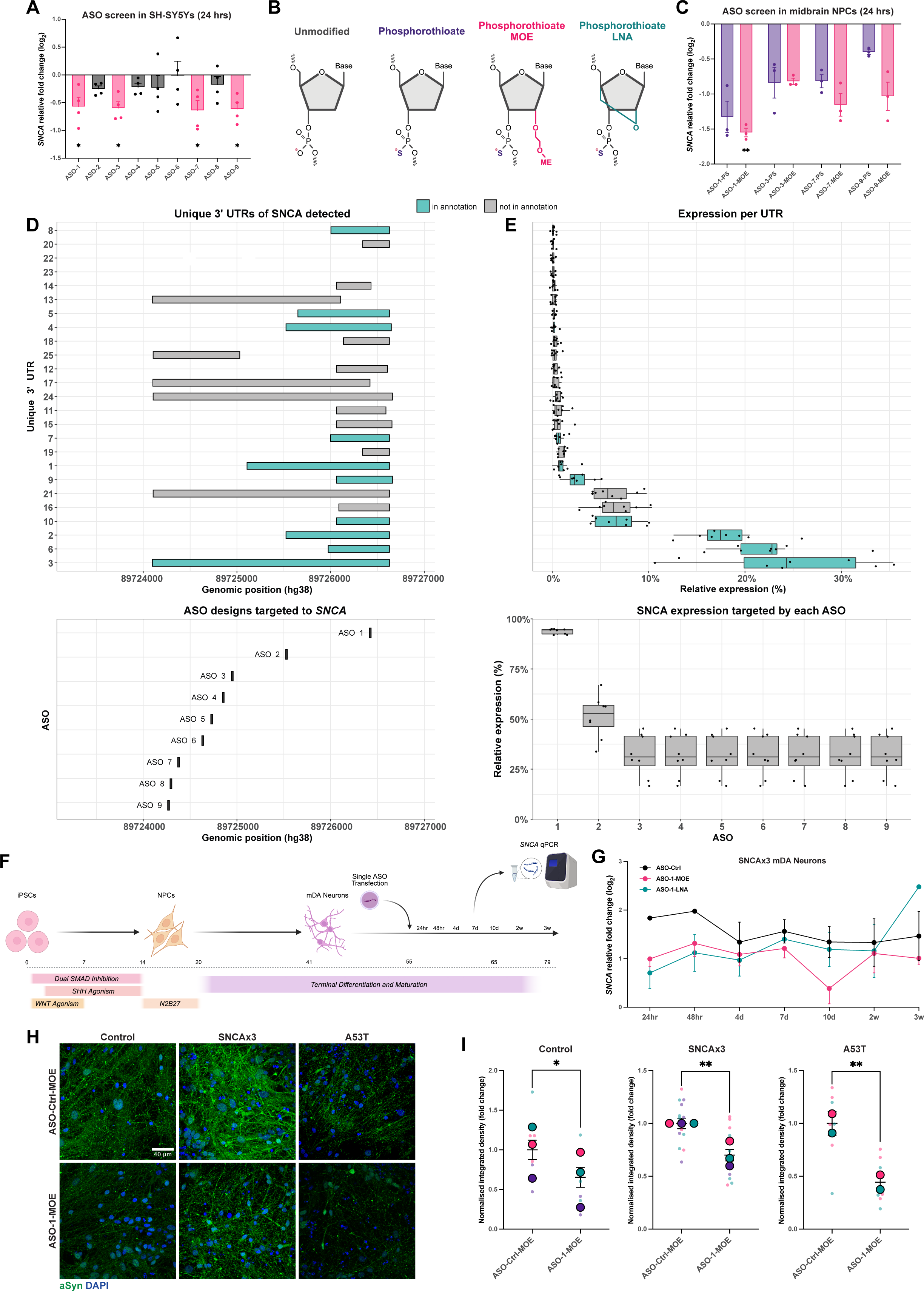
Design of ASOs targeted to *SNCA* transcripts. (**A**) Quantitative PCR showing *SNCA* mRNA expression in SH-SY5Ys treated with ASOs for 24 hours relative to a non-targeting control ASO (n = 4 independent experiments for each ASO, \**P* < 0.05, Kruskal-Wallis with Dunn’s multiple comparison test). (**B**) Graphic illustrating the chemical modifications used in the ASO designs. (**C**) Quantitative PCR showing *SNCA* mRNA expression in midbrain NPCs treated with ASOs for 24 hours relative to a non-targeting control ASO (n = 3 independent experiments for each ASO, \*\**P* < 0.01, Friedman’s test with Dunn’s multiple comparison test). (**D**) *SNCA* transcript structures detected in iPSC-derived mDA neurons and the targeting location of the 9 ASOs designed. (**E**) Relative *SNCA* expression by 3’UTR and the relative expression predicted to be targeted by each ASO sequence. (**F**) Schematic of the experimental design of treating mature mDA neurons with ASOs and quantifying *SNCA* expression at multiple timepoints after transfection. (**G**) Quantitative PCR data showing *SNCA* mRNA expression at multiple timepoints after ASO transfection in SNCAx3 iPSC-derived mDA neurons (n = 2 independent differentiations). (**H**) Representative ICC images showing the expression of total αSyn in control, SNCAx3, and A53T mDA neurons treated with a control and *SNCA*-targeting ASO for 10 days. (I) Quantification of the integrated fluorescence density of total αSyn in control, SNCAx3, and A53T mDA neurons with 10 days of ASO treatment (control = 3 donors, SNCAx3 = 3 independent differentiations, A53T = 2 donors; \**P* < 0.05, \*\**P* < 0.01, clustered Wilcoxon rank sum test).

The ASO chemistry alters ASO efficacy^35^ and therefore we tested the use of ASOs with sugar modifications (**Fig. 3B**) using a 5-10-5 gapmer design which retains the PS linkages throughout the ASO and has a central DNA gap (10mer) with the sugar modifications only for the bases on each flank. We modified the four ASOs that resulted in a significant knockdown in SH-SY5Ys after 24 hours to have the 2’-O-methoxyethyl (MOE) modification at the 2’ position. mDA NPCs were transfected with four ASOs +/-MOE gapmer design. qPCR analysis of *SNCA* in mDA NPCs 24 hours after transfection showed that for all ASOs, the MOE gapmer design achieved a greater knockdown relative to the respective ASO sequence with only a PS backbone.

Importantly, whilst all ASOs reduced the expression of *SNCA*, ASO-1 with the MOE modification (ASO-1-MOE) resulted in the greatest knockdown, and significantly reduced the levels of *SNCA* expression relative to ASO-Ctrl (**Fig. 3C**). Based on long-read RNAseq data ASO-1 would be expected to target the highest number of transcripts in neurons, accounting for over 93.8 ± 1.3% (range 92.0-95.0%) of transcription, whereas ASO-2 would target just 51.1 ± 10.7% (range 33.8-67.0%) of expression. The remainder of the ASOs only target 32.0 ± 10.6% (range 16.7-45.2%) of *SNCA* transcription (**Fig. 3D-E**).

ASO-1 was then selected for further modifications, specifically a locked nucleic acid (LNA) modification in a 5-10-5 gapmer design, where a methylene bridge connects the 2’-oxygen and the 4’-carbon of the ribose. We generated day 55 iPSC-derived mDA neurons and transfected them with ASO-1-PS, ASO-1-MOE, and ASO-1-LNA. ICC performed using an antibody that recognises total αSyn demonstrated that ASO-1-LNA resulted in a significant reduction in αSyn protein after 24 hours (**Fig. S8B-C**).

Next, we measured the persistence of the ASO effects in mDA neurons. Day 55 mDA control and SNCAx3 neurons were transfected with ASO-1-MOE and ASO-1-LNA and *SNCA* expression was measured by qPCR at 7 timepoints over a 3-week period (**Fig. 3F**). Consistent with our ICC analysis, in both genotypes there was a greater knockdown from the LNA modification at the earliest time point. However, the MOE modification resulted in a more sustained suppression of *SNCA*, with the greatest reduction in expression in SNCAx3 neurons occurring 10 days after transfection. Notably, in neurons treated with the LNA modification, a subsequent rebound increase in *SNCA* expression above the levels of those treated with the ASO-Ctrl occurred (**Fig. 3G, Fig. S8D**).

We further validated that ASO-1-MOE was specific for *SNCA*, demonstrating no significant difference in *SNCB* and *SNCG* expression in mDA neurons treated with ASO-1-MOE compared to ASO-Ctrl-MOE (**Fig. S9A-B**). We also confirmed that this ASO had no impact on neuronal enrichment through assessing the expression of key midbrain markers (*TH, SLC6A3, GIRK2, NURR1*; **Fig. S9C-F**). Finally, since ASO-1 was found to be targeting multiple *SNCA* transcript isoforms, we explored whether ASO treatment alters *SNCA* transcript usage in addition to overall gene expression (**Fig. S10-11**). We found that in ASO-1 treated SNCAx3 mDA neurons, αSyn-118 was no longer significantly upregulated compared to control neurons (**Fig. S11**) suggesting that ASO treatment aimed at reducing overall expression may also alter transcript usage.

We explored whether the reduction of *SNCA* expression after ASO-1-MOE treatment, and the alteration in *SNCA* transcript usage, was accompanied by reduced αSyn protein expression. Using an antibody to total αSyn, we show a significant decrease in αSyn as quantified by the integrated fluorescence density in control, SNCAx3, and A53T mDA neurons treated with ASO-1-MOE compared to ASO-NT-MOE (**Fig. 3H-I**). We also show a significant decrease in αSyn as quantified by the fluorescence area in SNCAx3 and A53T ASO-1-MOE treated mDA neurons (**Fig. S12B-C**).

### *SNCA* targeting ASO induces phenotypic reversal in patient mDA neurons

We previously reported the sequential development of cellular phenotypes in mDA neurons driven by the *SNCA* mutation.^31^ We investigated the effect of ASO-induced reduction in αSyn from day 55 to day 65 on these established αSyn-associated cellular phenotypes.

αSyn aggregation is a critical pathologic mechanism of PD,^4^ we therefore explored whether ASO treatment reduces pathologic neurotoxic forms of αSyn. Using a conformation-specific antibody to αSyn aggregates, we found a decrease in αSyn aggregation in SNCAx3 mutant mDA neurons treated with ASO-1-MOE based on the integrated fluorescence density (**Fig. 4A-B**) and fluorescence area (**Fig. S13A**) relative to ASO-Ctrl-MOE treated cells.

**Fig. 4.**
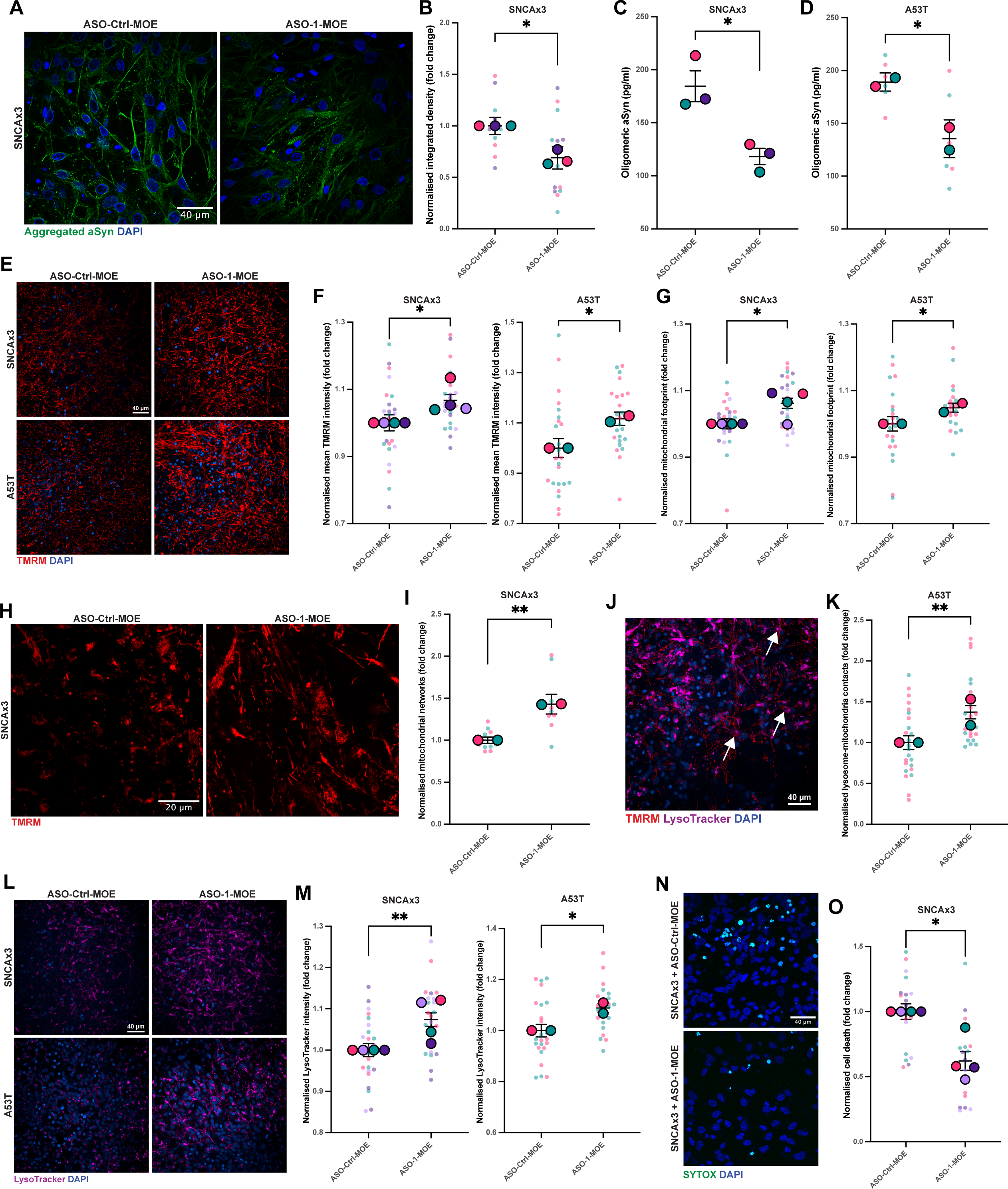
ASO-induced reduction of αSyn reduces PD-associated cellular phenotypes. (**A**) Representative ICC images showing aggregated αSyn using a conformation specific antibody in SNCAx3 mDA neurons with ASO treatment. (**B**) Quantification of the integrated fluorescence density of aggregated αSyn in ASO treated SNCAx3 mDA neurons (n = 3 independent differentiations; \**P* < 0.05, paired t-test). (**C**) Quantification of oligomeric αSyn by ELISA in SNCAx3 mDA neurons with ASO treatment (n = 3 independent differentiations; \**P* < 0.05, paired t-test). (**D**) Quantification of oligomeric αSyn by ELISA in A53T mDA neurons with ASO treatment (n = 2 donor lines each from 3 independent differentiations; \**P* < 0.05, clustered Wilcoxon rank sum test). (**E**) Representative live-cell high-throughput images of TMRM (mitochondria) in SNCAx3 and A53T mDA neurons treated with a control or *SNCA*-targeting ASO. (**F**) Quantification of the mean TMRM fluorescence intensity in SNCAx3 and A53T mDA neurons with ASO treatment (SNCAx3 = 4 independent differentiations, A53T = 2 lines across 4 independent differentiations; \**P* < 0.05, clustered Wilcoxon rank sum test). (**G**) Quantification of the mitochondrial footprint in SNCAx3 and A53T mDA neurons with ASO treatment (SNCAx3 = 4 independent differentiations, A53T = 2 lines across 4 independent differentiations; \**P* < 0.05, clustered Wilcoxon rank sum test). (**H**) Representative live-cell TMRM images in SNCAx3 mDA neurons treated with a control or *SNCA-*targeting ASO. (**I**) Assessment of the mitochondrial network in SNCAx3 mDA neurons with ASO treatment using MINA, \*\**P* < 0.01, paired t-test). (J) Representative live-cell high-throughput imaging of mitochondria (TMRM) and lysosome (LysoTracker Deep Red) contact sites in mDA neurons. (**K**) Quantification of the number of mitochondria-lysosome contacts in A53T mDA neurons treated with a control or *SNCA*-targeting ASO (A53T = 2 lines across 4 independent differentiations; \*\**P* < 0.01, clustered Wilcoxon rank sum test). (**L**) Representative live-cell high-throughput images of LysoTracker (lysosome) in SNCAx3 and A53T mDA neurons treated with a control or *SNCA*-targeting ASO. (**M**) Quantification of the mean LysoTracker fluorescence puncta intensity in SNCAx3 and A53T mDA neurons treated with a control or *SNCA*-targeting ASO (SNCAx3 = 4 independent experiments, A53T = 2 lines across 4 independent differentiations; \**P* < 0.05, \*\**P* < 0.01, clustered Wilcoxon rank sum test). (**N**) Live-cell images depicting dead cells in SNCAx3 mDA neurons with and without *SNCA*-targeting ASO treatment using the fluorescent dye SYTOX green. (**O**) Quantification of the proportion of dead cells in SNCAx3 mDA neurons with and without *SNCA*-targeting ASO treatment (SNCAx3 = 4 independent inductions; \**P* < 0.05, paired t-test).

An ELISA specific for oligomeric αSyn was used to determine the concentration of αSyn oligomers in the cells, and the concentration of secreted αSyn oligomers in the media and demonstrated a significantly reduced oligomer load in SNCAx3 and A53T mDA neurons treated with ASO-1-MOE compared to ASO-Ctrl-MOE (**Fig. 4C-D**). In A53T lines, there was a significant reduction in the secreted levels of αSyn oligomers, and non-significant reduction in the SNCAx3 neurons (**Fig. S13B-C**).

Mitochondrial dysfunction is an early critical pathway in PD pathogenesis,^36,37^ which is directly induced by αSyn aggregation^31,38,39^ and *SNCA* mDA neurons exhibit depolarised mitochondria at day 55. Using the lipophilic cationic dye TMRM as an indicator of mitochondrial membrane potential, in both the SNCAx3 line and in A53T mutant lines, ASO-1-MOE significantly increased (restored) mitochondrial membrane potential compared to treatment with ASO-Ctrl-MOE (**Fig. 4E-F**). ASO-1-MOE, compared to ASO-Ctrl-MOE, also significantly increased the mitochondrial footprint in mDA neurons (**Fig. 4G**), with less fragmentation of the mitochondrial network and a significant increase in the number of mitochondrial networks (**Fig. 4H-I**) and a non-significant increase in branch length (**Fig. S13D**). There was also a non-significant reduction in the number of donut-like mitochondria with ASO-1-MOE treatment (**Fig. S13E**), a mitochondrial morphology that is associated with mitochondrial stress.^40^

LysoTracker Deep Red, which accumulates in acidic cellular compartments, can be used in combination with TMRM to measure the polarised mitochondria-lysosome co-localisation, or contacts, in the cell. In ASO-1-MOE treated neurons, compared to ASO-Ctrl-MOE treated neurons, there was a significant increase in the number of contact sites between mitochondria and lysosomes in the A53T lines (**Fig. 4J-K**) and a non-significant increase in the SNCAx3 line **(Fig. S13F**). There was a significant increase in lysosome fluorescence intensity in ASO-1-MOE treated neurons compared to those treated with ASO-Ctrl-MOE in SNCAx3 and A53T neurons (**Fig. 4L-M)**. We previously demonstrated that textural features of both the lysosome and mitochondria are important in discerning disease states in PD iPSC-derived neurons.^41^ These features quantify spatial variation in pixel intensities in order to capture textural elements such as roughness or smoothness. Here, we showed that these features are significantly altered with ASO-1-MOE treatment compared to ASO-Ctrl-MOE for both the mitochondria and lysosome (**Fig. S13G-P**).

Finally, we explored whether an ASO-induced reduction in αSyn in mDA neurons, (once they had established alpha-synuclein pathology), altered SNCA induced toxicity. The fluorescent dye SYTOX-Green was used to identify the proportion of dead cells and demonstrated a significant reduction in cell death in SNCAx3 mDA neurons, and a non-significant reduction in A53T lines, treated with the ASO-1-MOE compared to ASO-Ctrl-MOE (**Fig. 4N-O, Fig. S13Q-R**).

## Discussion

The importance of *SNCA* transcript diversity, and the resulting αSyn isoforms, on PD pathogenicity remain poorly understood, despite being a likely fundamental determinant of the onset and progression of the disease. This study has unveiled 42 *SNCA* transcripts in iPSC-derived mDA neurons, a cell type vulnerable to degeneration in PD. We found no single dominant transcript, with the most highly expressed transcript accounting for less than 25% of total *SNCA* expression. Importantly, 12 of these transcripts were predicted to encode proteins with novel peptide sequences.

Previous research has focused exclusively on three shorter αSyn isoforms (αSyn-126, αSyn-112, and αSyn-98) demonstrating altered aggregation kinetics and lipid binding compared to the canonical αSyn-140,^15,16,42^ and brain region and disease dependent usage.^13,14,17,43–45^ Here, our targeted long-read data, derived from an enriched cell-type with high quality RNA, showed that about 10% of transcription is generated through previously unidentified protein-coding transcripts. Critically, we demonstrated that novel protein-coding transcripts are expressed across human brain regions obtained from adult post-mortem brain and are therefore not an artifact of an iPSC-derived model. Importantly, we showed that novel protein-coding isoforms detected in our iPSC-derived neurons are present in brain regions affected in PD and that some are translated (αSyn-128, αSyn-118 and αSyn-113). An *in-silico* analysis further demonstrated that compared to the canonical protein isoform (αSyn-140), these novel translated isoforms are predicted to have a lower solubility and are likely to be more aggregation prone. Although, we found both αSyn-118 and αSyn-113 at low levels compared to the canonical αSyn-140, it has been shown recently that small quantities of more aggregation prone αSyn isoforms increase the aggregation kinetics of αSyn-140.^16^ Therefore, our finding of novel aggregation-prone αSyn isoforms may have relevance to disease, despite their low relative abundance. We also found that αSyn-118 has a higher relative expression in iPSC mDA neurons derived from patients with *SNCA* mutation compared to those derived from healthy donors, further raising their possible relevance in contributing to disease pathogenesis.

However, it is important to note that much of the transcript diversity we identified at the *SNCA* locus was outside the ORF. In fact, we found that the majority of *SNCA* transcription in mDA neurons consisted of transcripts with a novel 3’ or 5’ UTRs (∼75%). Given that RNA transcript diversity regulates RNA localization, stability, and translational efficiency this is likely to be important.^46–49^ Given this, it is interesting that Rhinn *et al*. found enrichment of different 3’UTR lengths in PD patient brains and showed that alternative 3’UTR lengths impact the accumulation and localization (synaptic or mitochondrial) of αSyn,^22^ potentially through interactions with RNA-binding proteins.^50,51^ Similarly, there is evidence that αSyn regulation occurs through 5’UTR variability, with the presence of an iron responsive element,^52–54^ highlighting the possibility that translation of *SNCA* could be regulated by alternative 5’UTR usage and cellular iron. Together, these findings suggest that the robust identification of 5’ and 3’UTR variability in PD may reveal important insights into the pathophysiological processes driving disease.

These findings may also have important implications for the success of RNA targeting strategies in a clinical setting. Here, given the promise *SNCA* targeting ASOs have shown in animal models,^25–30^ we designed 9 ASOs targeting the 3’UTR of *SNCA* to explore the efficacy of this approach in a human patient cell model that faithfully recapitulates PD-associated cellular pathology. Targeted long-read analysis of *SNCA* demonstrates that due to the presence of a truncated 3’UTR in the majority of *SNCA* transcripts present in midbrain neurons, many ASO sequences designed against the canonical 3’UTR (the most unique region of *SNCA*) would only target a small proportion of total *SNCA* expression (∼30%). This observation highlights (i) the necessity of generating a comprehensive transcriptomic map when undertaking ASO design, and (ii) the importance of screening ASOs in a model with a transcriptional profile that reflects that of the target gene in the target cell.

Furthermore, whilst evaluation of ASO design is often focused on the likelihood of causing off-target effects, we demonstrated the importance of also considering unintended on-target effects. Targeted long-read RNA-seq of *SNCA* in ASO treated iPSC-derived mDA neurons illustrated that our *SNCA*-targeted ASO approach may modulate transcript usage, in addition to the intended total gene expression. This finding further emphasises the importance of screening gene manipulation strategies in a representative human model system, as it is likely that humanised animal models and non-human primates, that are primarily used for ASO development and validation, will have a different transcriptional and regulatory landscape.^55,56^

In fact, a more complete understanding of disease-relevant transcript structures may ultimately enable the generation of ASOs that specifically target disease-associated structures. In the context of *SNCA*, this more refined approach (rather than complete knockdown) may allow a higher level of *SNCA* transcription to be retained, which has important physiological functions,^57^ whilst still providing therapeutic benefit. Consistent with this view, we demonstrated that partial knockdown of *SNCA* is sufficient to elicit a phenotypic reversal, including reduced αSyn aggregation, amelioration of mitochondrial function, and improved cell viability. It should also be noted that a greater initial knockdown with ASO-1-LNA ultimately resulted in a significant upregulation in *SNCA* expression at subsequent timepoints, highlighting the importance of understanding the time-course and adaptation to this response.

Taken together, our study provides unique insights into the complexity of *SNCA* transcription and highlights its possible relevance to disease pathology. Our approach of combining a disease-relevant human patient neuronal model, targeted long-read RNA-seq, and ASO technology, revealed important considerations and complexities in the development of ASO gene manipulation strategies, highlighting the need for comprehensive transcriptomic mapping to inform therapeutic design. Overall, our research underscores the potential importance of developing nuanced approaches to *SNCA* manipulation, and strengthens the notion that *SNCA* targeting ASOs have the potential to be an effective disease-modifying therapy for PD.

## Acknowledgments

We would like to thank the patients for the fibroblast donation. We would like to thank all funding agencies. We would also like to thank the Francis Crick Institute Advanced Light Microscopy STP, the UCL Long Read Sequencing Service for their assistance and the UCL Biological Mass Spectrometry Centre. Some figure graphics were created with BioRender. This work is supported by the NIHR GOSH BRC. This research was funded in whole or in part by Aligning Science Across Parkinson’s [Grant numbers: ASAP-000509 (J.R.E., E.K.G., D.M., G.S.V., J.L., C.E.T., N.W.W., M.V., M.R., S.G.) and ASAP-000478 (H.M. and M.R.)] through the Michael J. Fox Foundation for Parkinson’s Research (MJFF), BrightFocus Foundation A2021009F (E.K.G.), Parkinson’s Disease UK G-2010 (W.H, K.M, and S.G) and Tenure Track Clinician Scientist Fellowship N008324/1 (M.R.). J.R.E. acknowledges support from the UCL Pat Harris PhD Fellowship in Neurodegenerative Disease. S.G. acknowledges funding from the i2i grant (The Francis Crick Institute), MJFF, the Wellcome Trust, and is an MRC Senior Clinical Fellow [MR/T008199/1]. The views expressed are those of the author(s) and not necessarily those of the NHS, the NIHR or the Department of Health. We are grateful to all members of the Ryten and Gandhi laboratories. For the purpose of open access, the author has applied a CC BY public copyright licence to all Author Accepted Manuscripts arising from this submission.

## Author contributions

Conceptualization: J.R.E., E.K.G., M.R. and S.G. Investigation: J.R.E., E.K.G., I.D., D.W., G.S.V., J.L., A.R., M.H.M., C.W.P., H.M., A.I.W., C.E.T., D.A., and M.L.C. Visualization: J.R.E. and E.K.G. Funding acquisition: N.W.W., M.V., M.R., and S.G. Project administration: J.R.E., E.K.G., M.R. and S.G. Supervision: N.W.W., M.V., K.M., W.H., M.R., and S.G. Writing – original draft: J.R.E., E.K.G., M.R. and S.G. Writing – review & editing: J.R.E., E.K.G., M.R. and S.G. All authors approved the final manuscript.

## STAR Methods

### Human induced pluripotent stem cell (hiPSC) maintenance

Parkinson’s patient and healthy control hiPSC lines (listed in **Supplementary Table 1**) were maintained in feeder-free monolayers on Geltrex™ (ThermoFisherScientific) coated 6-well plates and fed daily with mTeSR™1 (STEMCELL Technologies) medium. hiPSCs were passaged using 0.5 µM EDTA (Life Technologies), were maintained at 37 °C and 5% CO2, and underwent regular mycoplasma testing and short tandem repeat profiling.

### Midbrain dopaminergic neuron (mDA) differentiation

hiPSCs were grown to 100% and incubated with ‘N2B27’ medium and patterned for 14 days with daily media changes as previously described.^31^ After patterning, mDA neuronal precursor cells (NPCs) were maintained in ‘N2B27’ and were plated for terminal differentiation. Terminal differentiation medium consisted of ‘N2B27’ medium supplemented with 0.1 µM Compound E (Enzo Life Sciences) and 10 µM Rho kinase Rock inhibitor (Tocris).

### Antisense oligonucleotide synthesis

ASOs designed to target human *SNCA* and control ASOs with no targets in the human genome were synthesized and purified by IDT (Integrated DNA Technologies). ASO sequences are listed in **Supplementary Table 2**. ASOs were modified to have phosphorothioate linkages throughout. Where ASOs were modified with either MOE or LNA modifications a 5-10-5 gapmer design was used maintaining phosphorothioate linkages throughout.

### Antisense oligonucleotide transfection

Cells were transfected with *SNCA* targeting ASOs or a non-targeting control ASO with the same chemistry using DharmaFECT-1 transfection reagent (Horizon Discovery). The final concentration of ASO for the SH-SY5Y screen was 75 nm. For all experiments in iPSC-derived mDA NPCs or mDA neurons the final concentration was 300 nm.

### RNA extraction

Cell pellets were snap frozen using dry ice. RNA was harvested using the Maxwell^®^ RSC simplyRNA Cells kit (Promega) and the Maxwell^®^ RSC instrument or using the RNeasy kit followed by on-column DNase digestion (Qiagen) using the manufacturer’s protocols.

### Quantitative polymerase chain reaction

For qPCR, up to 2 µg of RNA was reverse-transcribed into cDNA using the High-Capacity cDNA Reverse Transcription kit (ThermoFisherScientific). qPCR was performed using the TaqMan^™^ Gene Expression Assay (ThermoFisherScientific) following the manufacturer’s protocol. TaqMan^™^ probes are listed in S**upplementary Table 3**. Samples were run for each gene in technical triplicate on the QuantStudio 6 Flex Real-Time PCR System (applied Biosystems). A minus reverse transcriptase control and no-template control were included as negative controls. Gene expression levels were normalized to the housekeeping genes, *GAPDH* and *ACTB*, following the delta-delta Ct method.

### Monomeric and Oligomeric ELISA

Monomeric ELISA was performed with the LEGEND MAX™ Human α-Synuclein ELISA Kit (SIG-38974, BioLegend) following manufacturer’s instructions. To determine oligomeric αSyn the Human Synuclein, alpha (non A4 component of amyloid precursor) oligomer (SNCA oligomer) ELISA kit (CSB-E18033h, Generon) was used following manufacturer’s instructions.

### Immunocytochemistry

Cells plated on Ibidi 8-well chambers were washed with PBS and fixed with 4% PFA for 15 minutes at RT. Samples were blocked and permeabilized in 5% BSA (Sigma) with 0.2% Triton-X-100 (Sigma) in PBS for 60 minutes. The cells were incubated with primary antibodies in 5% BSA at 4 °C overnight. Cells were then washed with PBS and stained with species-specific AlexaFluor conjugated secondary antibodies before being washed again with PBS and stained with Hoechst 33342 (ThermoFisherScientific). See **Supplementary Table 4** for all primary and secondary antibodies used and dilutions. Finally, cells were washed again with PBS before fluorescence mounting medium (Dako) was applied. Cells were stored at 4 °C until imaging. Image acquisition was conducted using the Zeiss 880 confocal system with a 40x 1.4 NA oil objective, and a pinhole of 1 AU. 5-6 Z projections were taken for each field of view.

### Live-cell imaging

Mitochondrial membrane potential, lysosomal dynamics, and cell death were measured using TMRM (25 nM), LysoTracker Deep Red (50 nM), SYTOX^™^ Green Nucleic Acid Stain (100 nM; all ThermoFisherScientific), and the nuclear marker Hoechst 3342. Cells were incubated with these dyes for 40 minutes in HBSS prior to imaging. Cells were either imaged using the Zeiss 880 confocal microscope with a 40x, 1.4 NA oil objective and a pinhole of 1 AU or were imaged using an Opera Phenix High-Content Screening System (PerkinElmer) as previously described.^41^ For both low and high-throughput imaging experiments 5-6 Z projections were taken for each field of view.

### Image analysis

ICC images and live-cell imaging obtained through the Zeiss 880 confocal microscope were analyzed using Fiji ImageJ. Fluorescent intensity, fluorescent area, and integrated density were measured across images with a threshold value set for all images in each dataset. To normalize the data, for each experiment, the fold change relative to the control condition was calculated. For mitochondria morphology analysis the ImageJ plugin “Mitochondrial Network Analysis (MiNA)” was used.^58^ For high-throughput live-cell imaging 20 fields of view were taken per well, data were collected by Columbus Studio Cell Analysis Software (Columbus 2.9.1, https://biii.eu/columbus-image-datastorage-and-analysis-system). All image analysis software used in this study are listed in **Supplementary Table 5**. Mitochondrial and lysosomal fluorescent features from high-throughput imaging were extracted using a previously validated pipeline.^41^ Detailed pipeline information can be found in **Supplementary Table 6**.

### Statistical analysis

Statistical analysis was either performed in Prism 10 or using R. Statistical tests for each experiment are specified in the figure legends. To account for clustered data (i.e. multiple cell, field of view, or well measurements within and across independent differentiations and donor lines), data were either averaged prior to statistical analysis or the Clustered Wilcoxon rank sum test using the Rosner-Glynn-Lee method was used to assess statistical significance between treatment groups. This was conducted in R using the “clusrank” package (https://cran.r-project.org/web/packages/clusrank/index.html).^59^ A *P* value below 0.05 was considered statistically significant. Results are represented as mean ± standard error of the mean (SEM). Where multiple sized dots are presented, large dots refer to the biological line (control and A53T) or to the independent differentiations where the experiment was conducted in a single donor line (SNCAx3). Small dots represent repeated measures, i.e. field of view or per well data, coloured by donor (control and A53T) or by independent differentiation (SNCAx3).

### CamSol and AlphaFold

Overall and residue-specific solubility scores of αSyn isoforms were calculated using the sequence-based CamSol Intrinsic pH-dependent software at pH 7.4. Isoelectric Point Calculator (IPC) 2.0 was used for pKa correction.^33,34^ Residue-specific solubility profiles were plotted using GraphPad Prism Version 10.2.0, with higher values indicating greater predicted solubility, and lower values indicating poorer predicted solubility, respectively.

To generate the conformational ensembles, AlphaFold ^60^ was used to predict the inter-residue distance distributions for each of the isoforms. These distance distributions were imposed as restraints in molecular dynamics simulations using metainference.^61^ The simulations were run via OpenMM and PLUMED^62^ using the coarse-grained CALVADOS-2^63^ force field. These simulations were run for up to 60 ns at pH 7.4 and 298 K. The figure shows the probability density plot of radius of gyration for each of the isoforms, overlaid with sample conformational ensembles coloured by their CamSol^33^ scores.

## PACBIO TARGETED ISO-SEQ

### cDNA synthesis

A total of 250ng of RNA was used per sample for reverse transcription. Two different cDNA synthesis approaches were used: (i) iPSC derived cell lines were generated using NEBNext® Single Cell/Low Input cDNA Synthesis & Amplification Module (New England Biolabs) and (ii) Human brain cDNA was generated by SMARTer PCR cDNA synthesis (Takara). For both reactions sample-specific barcoded oligo dT (12 µM) with PacBio 16mer barcode sequences were added (**Supplementary Table 7**).

#### SMARTer PCR cDNA synthesis

First strand synthesis was performed as per manufacturer instructions, using sample-specific barcoded primers instead of the 3’ SMART CDS Primer II A. We used a 90 min incubation to generate full-length cDNAs. cDNA amplification was performed using a single primer (5’ PCR Primer II A from the SMARTer kit, 5′ AAG CAG TGG TAT CAA CGC AGA GTA C 3′) and was used for all PCR reactions post reverse transcription. We followed the manufacturer’s protocol with our determined optimal number of 18 cycles for amplification; this was used for all samples. We used a 6 min extension time in order to capture longer cDNA transcripts. PCR products were purified separately with 1X ProNex® Beads.

#### NEBNext® Single Cell/Low Input cDNA Synthesis & Amplification Module

A reaction mix of 5.4 μL of total RNA (250 ng in total), 2 μL of barcoded primer, 1.6 μL of dNTP (25 mM) held at 70°C for 5 min. This reaction mix was then combined with 5 μL of NEBNext Single Cell RT Buffer, 3 μL of nuclease-free H_2_O and 2 μL NEBNext Single Cell RT Enzyme Mix. The reverse transcription mix was then placed in a thermocycler at 42°C with the lid at 52°C for 75 minutes then held at 4°C. On ice, we added 1 μL of Iso-Seq Express Template Switching Oligo and then placed the reaction mix in a thermocycler at 42°C with the lid at 52°C for 15 minutes. We then added 30 μL elution buffer (EB) to the 20 μL Reverse Transcription and Template Switching reaction (for a total of 50 μL), which was then purified with 1X ProNex® Beads and eluted in 46 μL of EB. cDNA amplification was performed by combining the eluted Reverse Transcription and Template Switching reaction with 50 μL of NEBNext Single Cell cDNA PCR Master Mix, 2 μL of NEBNext Single Cell cDNA PCR Primer, 2 μL of Iso-Seq Express cDNA PCR Primer and 0.5 μL of NEBNext Cell Lysis Buffer.

### cDNA Capture Using IDT Xgen® Lockdown® Probes

We used the xGen Hyb Panel Design Tool (https://eu.idtdna.com/site/order/designtool/index/XGENDESIGN) to design non-overlapping 120-mer hybridization probes against *SNCA*. We removed any overlapping probes with repetitive sequences (repeatmasker) and to reduce the density of probes mapping to intronic regions 0.2, which means 1 probe per 1.2kb. In the end, our probe pool consisted of 111 probes targeted towards *SNCA* (**Fig. S6**).

We pooled an equal mass of barcoded cDNA for a total of 500 ng per capture reaction. Pooled cDNA was combined with 7.5 μL of Cot DNA in a 1.5 mL LoBind tube. We then added 1.8X of ProNex beads to the cDNA pool with Cot DNA, gently mixed the reaction mi× 10 times (using a pipette) and incubated for 10 min at room temperature. After two washes with 200 μL of freshly prepared 80% ethanol, we removed any residual ethanol and immediately added 19 μL hybridization mix consisting of: 9.5 μL of 2X Hybridization Buffer, 3 μL of Hybridization Buffer Enhancer, 1 μL of xGen Asym TSO block (25 nmole), 1 μL of polyT block (25 nmole) and 4.5 μL of 1X xGen Lockdown Probe pool.

The PacBio targeted Iso-Seq protocol is described in detail at protocols.io (dx.doi.org/10.17504/protocols.io.n92ld9wy9g5b/v1).

### Automated Analysis of Iso-Seq data using Snakemake

For the analysis of targeted PacBio Iso-Seq data, we created two Snakemake^64^ (v 5.32.2; RRID:SCR_003475) pipelines to analyse targeted long-read RNA-seq robustly and systematically:

#### APTARS

(Analysis of PacBio TARgeted Sequencing, https://github.com/sid-sethi/APTARS): For each SMRT cell, two files were required for processing: (i) a subreads.bam and (ii) a FASTA file with primer sequences, including barcode sequences.

Each sequencing run was processed by ccs (v 5.0.0; RRID:SCR_021174; https://ccs.how/), which combines multiple subreads of the same SMRTbell molecule and to produce one highly accurate consensus sequence, also called a HiFi read (≥ Q20). We used the following parameters: -- minLength 10 –maxLength 50000 –minPasses 3 –minSnr 2.5 –maxPoaCoverage 0 – minPredictedAccuracy 0.99.

Identification of barcodes, demultiplexing and removal of primers was then performed using lima (v 2.0.0; https://lima.how/) invoking –isoseq –peek-guess.

Isoseq3 (v 3.4.0; https://github.com/PacificBiosciences/IsoSeq) was then used to (i) remove polyA tails and (ii) identify and remove concatemers using, with the following parameters refine – require-polya, --log-level DEBUG. This was followed by clustering and polishing with the following parameters using: cluster flnc.fofn clustered.bam –verbose –use-qvs.

Reads with predicted accuracy ≥ 0.99 were aligned to the GRCh38 reference genome using minimap2^65^ (v 2.17; RRID:SCR_018550) using -ax splice:hq -uf –secondary=no. samtools^66^ (RRID:SCR_002105; http://www.htslib.org/) was then used to sort and filter the output SAM for the locus of gene of interest, as defined in the config.yml.

We used cDNA_Cupcake (v 22.0.0; https://github.com/Magdoll/cDNA_Cupcake) to: (i) collapse redundant transcripts, using collapse_isoforms_by_sam.py (--dun-merge-5-shorter) and (ii) obtain read counts per sample, using get_abundance_post_collapse.py followed by demux_isoseq_with_genome.py.

Isoforms detected were characterized and classified using SQANTI3^67^ (v 4.2; https://github.com/ConesaLab/SQANTI3) in combination with GENCODE (v 38) comprehensive gene annotation. An isoform was classified as full splice match (FSM) if it aligned with reference genome with the same splice junctions and contained the same number of exons, incomplete splice match (ISM) if it contained fewer 5′ exons than reference genome, novel in catalog (NIC) if it is a novel isoform containing a combination of known donor or acceptor sites, or novel not in catalog (NNC) if it is a novel isoform with at least one novel donor or acceptor site.

#### PSQAN

(Post Sqanti QC Analysis, https://github.com/sid-sethi/PSQAN) Following transcript characterisation from SQANTI3, we applied a set of filtering criteria to remove potential genomic contamination and rare PCR artifacts. We removed an isoform if: (1) the percent of genomic “A” s in the downstream 20 bp window was more than 80% (“perc_A_downstream_TTS” > 80); (2) one of the junctions was predicted to be template switching artifact (“RTS_stage” = TRUE); or (3) it was not associated with the gene of interest. Using SQANTI’s output of ORF prediction, NMD prediction and structural categorisation based on comparison with the reference annotation (GENCODE), we grouped the identified isoforms into the following categories: **(1) Non-coding novel –** if predicted to be non-coding and not a full-splice match with the reference; **(2) Non-coding known –** if predicted to be non-coding and a full-splice match with the reference; **(3) NMD novel –** if predicted to be coding & NMD, and not a full-splice match with the reference; **(4) NMD known –** if predicted to be coding & NMD, and a full-splice match with the reference; **(5) Coding novel –** if predicted to be coding & not NMD, and not a full-splice match with the reference; **(6) Coding known (complete match)** – if predicted to be coding & not NMD, and a full-splice & UTR match with the reference; and **(7) Coding known (alternate 3’/5’ end) –** if predicted to be coding & not NMD, and a full-splice match with the reference but with an alternate 3’ end, 5’ end or both 3’ and 5’ end.

Given a transcript *T* in sample ” with *FLR* as the number of full-length reads mapped to the transcript *T*, we calculated the normalised full-length reads (*NFLR_Ti_*) as the percentage of total transcription in the sample:

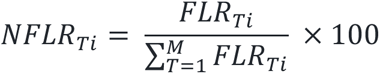

where, *NFLR_Ti_* represents the normalised full-length read count of transcript *T* in sample *i*, *FLR_Ti_* is the full-length read count of transcript *T* in sample *i* and *M* is the total number of transcripts identified to be associated with the gene after filtering. Finally, to summarise the expression of a transcript associated with a gene, we calculated the mean of normalised full-length reads (*NFLR_Ti_*) across all the samples:

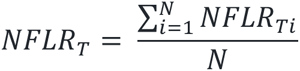

where, *NFLR_T_* represents the mean expression of transcript *T* across all samples and *N* is the total number of samples. To remove low-confidence isoforms arising from artefacts, we only selected isoforms fulfilling the following three criteria: (1) expression of m inimum 0.1% of total transcription per sample, i.e., *NFLR_Ti_* ≥ 0.1; (2) a minimum of 80% of total samples passing the *NFLR_Ti_* threshold; and (3) expression of minimum 0.3% of total transcription across samples, i.e., *NFLR_T_* ≥ 0.3.

### Visualizations of transcripts

For any visualization of transcript structures, we have recently developed ggtranscript ^68^ (v 0.99.03; https://github.com/dzhang32/ggtranscript), a R package that extends the incredibly popular tool ggplot2 (v 3.3.5 RRID; SCR_014601) for visualizing transcript structure and annotation.

### Proteomic analysis of isoforms using TX2P – Transcript to protein

We have developed a tool TX2P - Transcript to protein which allows for automated integration of mass spectometry data in long-read RNA-seq workflows (https://github.com/MurphyDavid/TX2P.git). It does so by predicting open reading frames for each transcript within an input GTF/GFF file, translates it into peptide sequences and then searches for those predicted proteins in mass spec datasets using MetaMorpheus^69^ (v 0.0.320; https://github.com/smith-chem-wisc/MetaMorpheus). A total of 30 public mass spectrometry data was retrieved from ProteomeXchange and MassIVE (**Supplementary Table 8**).

### Open reading frame expression in iPSC and human brain

To analyse the expression of open reading frames (ORFs), we merged transcripts sharing identical ORFs, irrespective of their 5’ and 3’ UTR structures. Subsequently, we normalized the expression levels, focusing solely on these merged transcripts.

## ALPHA-SYNUCLEIN ISOFORM MASS SPECTROMETRY

### Tissue preparation

Post-mortem human brain tissue was obtained from the Queen Square Brain Bank, University College London. Informed consent was given in all cases (**Supplementary Table 9**). Control subjects did not have a diagnosis of neurological disease in life and at post-mortem displayed age-related changes only. All cases were assessed for αSyn pathology and rated according to Braak staging. Ethical approval for the study was obtained from the Local Research Ethics Committee of the National Hospital for Neurology and Neurosurgery. Frozen post-mortem human brain samples from the anterior cingulate cortex and substantia nigra were homogenised in 1ml RIPA buffer (Pierce) with phosphatase inhibitors (Roche) and protease inhibitors (Roche) using Kimble dounce tissue grinders (D8938) for a total of 50 douncing cycles, 25 x A and 25 x B. Lysates were centrifuged at 4000rpm for 10min at 4°C and 1ml supernatant collected for immunoprecipitation. Additional volumes were collected for further validation.

### Immunoprecipitation

The recombinant αSyn antibody (MJFR-1) was centrifuged at 16000 x g for 10 minutes at 4°C. Antibody coupling kit M-270 Epoxy Dynabeads (14311D, Thermofisher Scientific) were covalently coupled to MJFR-1 αSyn antibody at a concentration of 7μg antibody/mg bead and stored in a final bead concentration of 10mg/ml antibody-coupled beads in SB buffer (following manufacturer’s protocol).

Dynabeads were redistributed by vortex and 500μl of beads were added to each lysate and incubated for 24 hours at room temperature on a constant roller. Magnets were used to collect bead pellets and the supernatant was saved. Pellets were stored at -20°C prior to direct elution.

### Protein Preparation and Digestion

Proteins immobilized on magnetic beads post-immunoprecipitation were processed as follows: the beads were thrice washed with Phosphate Buffered Saline (PBS) utilizing a magnetic separation rack to eliminate residual buffers. Subsequently, beads were resuspended in 20 µL of digestion buffer (6M Urea, 2M Thiourea, 2% ASB-14 in 200 mM Tris-HCl, pH 8.0) and incubated at room temperature for 1 hour with agitation to denature proteins. Reduction of disulfide bonds was achieved by the addition of 3 µL of 169 mM Dithiothreitol (DTT) and incubation for 1 hour at room temperature with shaking. Alkylation was performed by adding 6 µL of 169 mM Iodoacetamide (IAA) and incubating for 45 minutes in the dark at room temperature. To facilitate efficient trypsin digestion, the mixture was diluted with 165 µL of Milli-Q water to decrease the urea concentration below 1M. Trypsin Gold was prepared at a concentration of 0.1 µg/µL in the manufacturer’s provided resuspension buffer. Digestion was initiated by adding 10 µL of the trypsin solution (1 µg) to each sample, followed by brief vortexing. The samples were then digested for 10 hours at 37°C with shaking in a thermomixer. The reaction was quenched by the addition of 200 µL of 0.2% Trifluoroacetic Acid (TFA).

### Peptide Cleanup via Solid Phase Extraction (SPE)

Following trypsin digestion, peptide mixtures were subjected to a purification process using Solid Phase Extraction (SPE). SPE cartridges (Biotage, Sweden), which were pre-wetted twice with 1 mL of 60% Acetonitrile (ACN) in 0.1% Trifluoroacetic Acid (TFA). This was followed by an equilibration step, where cartridges were treated twice with 1 mL of 0.1% TFA. Acidified peptide samples were then loaded onto the equilibrated cartridges and allowed to drip through under gravity. To further purify captured peptides, the cartridges were washed twice with 0.1% TFA. Peptides were subsequently eluted into 1.5 mL microtubes using two sequential aliquots of 500 µL of 60% ACN in 0.1% TFA. The eluted peptides were then dried using a rotational evaporator under vacuum at room temperature for 7.5 hours, resulting in dry peptide samples that were stored at - 20⁰ C until analysis.

### Assay Development and Library Construction

In the investigation of novel alpha-synuclein isoforms, sequences were aligned, and unique tryptic peptides were identified. These peptides were custom synthesised (Genscript USA) to optimise the method. Peptide sequences were imported into Skyline (Macoss Labs USA) a freely-available, open-source Windows client application for building Multiple Reaction Monitoring (MRM) methods. In conjunction with Prosit, an in-silico spectral library was generated for each peptide across different precursor charge states (+2, +3, and +4). A normalized collision energy (NCE) of 31, experimentally established for this instrument using a synthetic peptide pool, was utilized for library creation. Fragment selection for the specified charge states was refined by filtering to retain the top 12 most intense fragments predicted by the in-silico spectral library. This process facilitated the export of multiple acquisition methods to MassLynx v 4.2 (Waters Corp USA), each configured with a minimum dwell time per transition of 10 microseconds. A peptide unique to the rabbit antibody used to immunoprecipitate the αSyn was included in the assay to use for normalisation of total αSyn. Transitions used in the targeted assay are listed in **Supplementary Table 10**.

### Sample Preparation and Chromatographic Conditions

Dried peptide digests were reconstituted in 100 µL of 5% ACN and 0.1% TFA. Chromatographic separation was achieved on a 10 cm Aquity Premier column (2.1mm diameter, 1.7 µm particle size), maintaining the column at 60°C. The injection volume was 5 µL. Initial mobile phase conditions were set at 95% A (0.1% Formic acid in water) and 5% B (0.1% Formic acid in acetonitrile), held for 1 minute before applying a gradient over 13.4 minutes to reach 35% B, followed by a ramp to 100% B 1 minute, and maintained for an additional 2 minutes. The system was then returned to initial conditions for column equilibration, with a total run time of 20 minutes per injection.

### Instrument Parameters and Data Acquisition

Instrumentation was operated in positive ion mode, with a capillary voltage of 2.8kV and a cone voltage maintained at 35V. The source and desolvation temperatures were set at 150°C and 500°C, respectively, with desolvation and cone gas flows at 700 L/hour and 150 L/hour. Following data acquisition, raw data were imported into Skyline for evaluation, with peptide identity confirmed through spectral library matching (dot product score cutoff of 0.75), matching a minimum of 5 transitions (**Fig. S6-7**). The selection of predominant precursor charges and fragments was based on intensity and chromatographic performance, facilitating another round of method export focusing on collision energy optimization for selected precursors and fragments. This final optimised method was applied to samples and a pooled sample, comprising 30 µL from each digest, was used to QC the analysis. Results were exported to a CSV file and only peak data with <20% CV were accepted for further analysis in excel.

### Data and code availability

Raw long-read RNAseq data generated and utilized in this manuscript are publicly available in Synapse (Project SynID: syn53642785): https://www.synapse.org/#!Synapse:syn53642785/wiki/626685. The code for analysis and figure generation in this manuscript can be accessed through: https://github.com/egustavsson/SNCA_manuscript.git.

## List of Supplementary Materials

Fig S1 to S13

Tables S1 to S12

## References

1. Shahmoradian, S.H., Lewis, A.J., Genoud, C., Hench, J., Moors, T.E., Navarro, P.P., Castaño-Díez, D., Schweighauser, G., Graff-Meyer, A., Goldie, K.N., et al. (2019). Lewy pathology in Parkinson’s disease consists of crowded organelles and lipid membranes. Nat. Neurosci. 22, 1099–1109. 10.1038/s41593-019-0423-2.

2. McCann, H., Stevens, C.H., Cartwright, H., and Halliday, G.M. (2014). α-Synucleinopathy phenotypes. Parkinsonism Relat. Disord. 20 *Suppl 1*, S62–7. 10.1016/S1353-8020(13)70017-8.

3. Cookson, M.R. (2009). alpha-Synuclein and neuronal cell death. Mol. Neurodegener. 4, 9. 10.1186/1750-1326-4-9.

4. Choi, M.L., and Gandhi, S. (2018). Crucial role of protein oligomerization in the pathogenesis of Alzheimer’s and Parkinson’s diseases. FEBS J. 285, 3631–3644. 10.1111/febs.14587.

5. Singleton, A.B., Farrer, M., Johnson, J., Singleton, A., Hague, S., Kachergus, J., Hulihan, M., Peuralinna, T., Dutra, A., Nussbaum, R., et al. (2003). alpha-Synuclein locus triplication causes Parkinson’s disease. Science 302, 841. 10.1126/science.1090278.

6. Book, A., Guella, I., Candido, T., Brice, A., Hattori, N., Jeon, B., Farrer, M.J., and SNCA Multiplication Investigators of the GEoPD Consortium (2018). A Meta-Analysis of α-Synuclein Multiplication in Familial Parkinsonism. Front. Neurol. 9, 1021. 10.3389/fneur.2018.01021.

7. Nishioka, K., Ross, O.A., Ishii, K., Kachergus, J.M., Ishiwata, K., Kitagawa, M., Kono, S., Obi, T., Mizoguchi, K., Inoue, Y., et al. (2009). Expanding the clinical phenotype of SNCA duplication carriers. Mov. Disord. 24, 1811–1819. 10.1002/mds.22682.

8. Konno, T., Ross, O.A., Puschmann, A., Dickson, D.W., and Wszolek, Z.K. (2016). Autosomal dominant Parkinson’s disease caused by SNCA duplications. Parkinsonism Relat. Disord. 22 *Suppl 1*, S1–6. 10.1016/j.parkreldis.2015.09.007.

9. Mata, I.F., Shi, M., Agarwal, P., Chung, K.A., Edwards, K.L., Factor, S.A., Galasko, D.R., Ginghina, C., Griffith, A., Higgins, D.S., et al. (2010). SNCA variant associated with Parkinson disease and plasma alpha-synuclein level. Arch. Neurol. 67, 1350–1356. 10.1001/archneurol.2010.279.

10. Campêlo, C.L. das C., and Silva, R.H. (2017). Genetic variants in SNCA and the risk of sporadic parkinson’s disease and clinical outcomes: A review. Parkinsons. Dis. 2017, 4318416. 10.1155/2017/4318416.

11. Polymeropoulos, M.H., Lavedan, C., Leroy, E., Ide, S.E., Dehejia, A., Dutra, A., Pike, B., Root, H., Rubenstein, J., Boyer, R., et al. (1997). Mutation in the alpha-synuclein gene identified in families with Parkinson’s disease. Science 276, 2045–2047. 10.1126/science.276.5321.2045.

12. Papadimitriou, D., Antonelou, R., Miligkos, M., Maniati, M., Papagiannakis, N., Bostantjopoulou, S., Leonardos, A., Koros, C., Simitsi, A., Papageorgiou, S.G., et al. (2016). Motor and Nonmotor Features of Carriers of the p.A53T Alpha-Synuclein Mutation: A Longitudinal Study. Mov. Disord. 31, 1226–1230. 10.1002/mds.26615.

13. Beyer, K., Domingo-Sábat, M., Lao, J.I., Carrato, C., Ferrer, I., and Ariza, A. (2008). Identification and characterization of a new alpha-synuclein isoform and its role in Lewy body diseases. Neurogenetics 9, 15–23. 10.1007/s10048-007-0106-0.

14. Beyer, K., Humbert, J., Ferrer, A., Lao, J.I., Carrato, C., López, D., Ferrer, I., and Ariza, A. (2006). Low alpha-synuclein 126 mRNA levels in dementia with Lewy bodies and Alzheimer disease. Neuroreport 17, 1327–1330. 10.1097/01.wnr.0000224773.66904.e7.

15. Bungeroth, M., Appenzeller, S., Regulin, A., Völker, W., Lorenzen, I., Grötzinger, J., Pendziwiat, M., and Kuhlenbäumer, G. (2014). Differential aggregation properties of alpha-synuclein isoforms. Neurobiol. Aging 35, 1913–1919. 10.1016/j.neurobiolaging.2014.02.009.

16. Röntgen, A., Toprakcioglu, Z., Tomkins, J.E., and Vendruscolo, M. (2024). Modulation of α-synuclein in vitro aggregation kinetics by its alternative splice isoforms. Proc. Natl. Acad. Sci. USA 121, e2313465121. 10.1073/pnas.2313465121.

17. Beyer, K., Lao, J.I., Carrato, C., Mate, J.L., López, D., Ferrer, I., and Ariza, A. (2004). Differential expression of alpha-synuclein isoforms in dementia with Lewy bodies. Neuropathol Appl Neurobiol 30, 601–607. 10.1111/j.1365-2990.2004.00572.x.

18. Kalivendi, S.V., Yedlapudi, D., Hillard, C.J., and Kalyanaraman, B. (2010). Oxidants induce alternative splicing of alpha-synuclein: Implications for Parkinson’s disease. Free Radic. Biol. Med. 48, 377–383. 10.1016/j.freeradbiomed.2009.10.045.

19. Mitschka, S., and Mayr, C. (2022). Context-specific regulation and function of mRNA alternative polyadenylation. Nat. Rev. Mol. Cell Biol. 23, 779–796. 10.1038/s41580-022-00507-5.

20. Doxakis, E. (2010). Post-transcriptional regulation of alpha-synuclein expression by mir-7 and mir-153. J. Biol. Chem. 285, 12726–12734. 10.1074/jbc.M109.086827.

21. Junn, E., Lee, K.-W., Jeong, B.S., Chan, T.W., Im, J.-Y., and Mouradian, M.M. (2009). Repression of alpha-synuclein expression and toxicity by microRNA-7. Proc. Natl. Acad. Sci. USA 106, 13052–13057. 10.1073/pnas.0906277106.

22. Rhinn, H., Qiang, L., Yamashita, T., Rhee, D., Zolin, A., Vanti, W., and Abeliovich, A. (2012). Alternative α-synuclein transcript usage as a convergent mechanism in Parkinson’s disease pathology. Nat. Commun. 3, 1084. 10.1038/ncomms2032.

23. Tseng, E., Rowell, W.J., Glenn, O.-C., Hon, T., Barrera, J., Kujawa, S., and Chiba-Falek, O. (2019). The landscape of SNCA transcripts across synucleinopathies: new insights from long reads sequencing analysis. Front. Genet. 10, 584. 10.3389/fgene.2019.00584.

24. Bennett, C.F., Krainer, A.R., and Cleveland, D.W. (2019). Antisense oligonucleotide therapies for neurodegenerative diseases. Annu. Rev. Neurosci. 42, 385–406. 10.1146/annurev-neuro-070918-050501.

25. Alarcón-Arís, D., Pavia-Collado, R., Miquel-Rio, L., Coppola-Segovia, V., Ferrés-Coy, A., Ruiz-Bronchal, E., Galofré, M., Paz, V., Campa, L., Revilla, R., et al. (2020). Anti-α-synuclein ASO delivered to monoamine neurons prevents α-synuclein accumulation in a Parkinson’s disease-like mouse model and in monkeys. EBioMedicine 59, 102944. 10.1016/j.ebiom.2020.102944.

26. Pavia-Collado, R., Cóppola-Segovia, V., Miquel-Rio, L., Alarcón-Aris, D., Rodríguez-Aller, R., Torres-López, M., Paz, V., Ruiz-Bronchal, E., Campa, L., Artigas, F., et al. (2021). Intracerebral Administration of a Ligand-ASO Conjugate Selectively Reduces α-Synuclein Accumulation in Monoamine Neurons of Double Mutant Human A30P*A53T*α-Synuclein Transgenic Mice. Int. J. Mol. Sci. 22. 10.3390/ijms22062939.

27. Uehara, T., Choong, C.-J., Nakamori, M., Hayakawa, H., Nishiyama, K., Kasahara, Y., Baba, K., Nagata, T., Yokota, T., Tsuda, H., et al. (2019). Amido-bridged nucleic acid (AmNA)-modified antisense oligonucleotides targeting α-synuclein as a novel therapy for Parkinson’s disease. Sci. Rep. 9, 7567. 10.1038/s41598-019-43772-9.

28. Brown, J.L., Hart, D.W., Boyle, G.E., Brown, T.G., LaCroix, M., Baraibar, A.M., Pelzel, R., Kim, M., Sherman, M.A., Boes, S., et al. (2022). SNCA genetic lowering reveals differential cognitive function of alpha-synuclein dependent on sex. Acta Neuropathol. Commun. 10, 180. 10.1186/s40478-022-01480-y.

29. Cole, T.A., Zhao, H., Collier, T.J., Sandoval, I., Sortwell, C.E., Steece-Collier, K., Daley, B.F., Booms, A., Lipton, J., Welch, M., et al. (2021). α-Synuclein antisense oligonucleotides as a disease-modifying therapy for Parkinson’s disease. JCI Insight 6. 10.1172/jci.insight.135633.

30. Yang, J., Luo, S., Zhang, J., Yu, T., Fu, Z., Zheng, Y., Xu, X., Liu, C., Fan, M., and Zhang, Z. (2021). Exosome-mediated delivery of antisense oligonucleotides targeting α-synuclein ameliorates the pathology in a mouse model of Parkinson’s disease. Neurobiol. Dis. 148, 105218. 10.1016/j.nbd.2020.105218.

31. Virdi, G.S., Choi, M.L., Evans, J.R., Yao, Z., Athauda, D., Strohbuecker, S., Nirujogi, R.S., Wernick, A.I., Pelegrina-Hidalgo, N., Leighton, C., et al. (2022). Protein aggregation and calcium dysregulation are hallmarks of familial Parkinson’s disease in midbrain dopaminergic neurons. npj Parkinsons Disease 8, 162. 10.1038/s41531-022-00423-7.

32. Rydbirk, R., Østergaard, O., Folke, J., Hempel, C., DellaValle, B., Andresen, T.L., Løkkegaard, A., Hejl, A.-M., Bode, M., Blaabjerg, M., et al. (2022). Brain proteome profiling implicates the complement and coagulation cascade in multiple system atrophy brain pathology. Cell Mol. Life Sci. 79, 336. 10.1007/s00018-022-04378-z.

33. Sormanni, P., Aprile, F.A., and Vendruscolo, M. (2015). The CamSol method of rational design of protein mutants with enhanced solubility. J. Mol. Biol. 427, 478–490. 10.1016/j.jmb.2014.09.026.

34. Oeller, M., Kang, R., Bell, R., Ausserwöger, H., Sormanni, P., and Vendruscolo, M. (2023). Sequence-based prediction of pH-dependent protein solubility using CamSol. Brief. Bioinformatics 24. 10.1093/bib/bbad004.

35. Crooke, S.T., Vickers, T.A., and Liang, X.-H. (2020). Phosphorothioate modified oligonucleotide-protein interactions. Nucleic Acids Res. 48, 5235–5253. 10.1093/nar/gkaa299.

36. Bose, A., and Beal, M.F. (2016). Mitochondrial dysfunction in Parkinson’s disease. J. Neurochem. 139 *Suppl 1*, 216–231. 10.1111/jnc.13731.

37. Toomey, C.E., Heywood, W.E., Evans, J.R., Lachica, J., Pressey, S.N., Foti, S.C., Al Shahrani, M., D’Sa, K., Hargreaves, I.P., Heales, S., et al. (2022). Mitochondrial dysfunction is a key pathological driver of early stage Parkinson’s. Acta Neuropathol. Commun. 10, 134. 10.1186/s40478-022-01424-6.

38. Choi, M.L., Chappard, A., Singh, B.P., Maclachlan, C., Rodrigues, M., Fedotova, E.I., Berezhnov, A.V., De, S., Peddie, C.J., Athauda, D., et al. (2022). Pathological structural conversion of α-synuclein at the mitochondria induces neuronal toxicity. Nat. Neurosci. 25, 1134–1148. 10.1038/s41593-022-01140-3.

39. Ludtmann, M.H.R., Angelova, P.R., Horrocks, M.H., Choi, M.L., Rodrigues, M., Baev, A.Y., Berezhnov, A.V., Yao, Z., Little, D., Banushi, B., et al. (2018). α-synuclein oligomers interact with ATP synthase and open the permeability transition pore in Parkinson’s disease. Nat. Commun. 9, 2293. 10.1038/s41467-018-04422-2.

40. Picard, M., and McEwen, B.S. (2014). Mitochondria impact brain function and cognition. Proc. Natl. Acad. Sci. USA 111, 7–8. 10.1073/pnas.1321881111.

41. D’Sa, K., Evans, J.R., Virdi, G.S., Vecchi, G., Adam, A., Bertolli, O., Fleming, J., Chang, H., Leighton, C., Horrocks, M.H., et al. (2023). Prediction of mechanistic subtypes of Parkinson’s using patient-derived stem cell models. Nat. Mach. Intell. 5, 933–946. 10.1038/s42256-023-00702-9.

42. Soll, L.G., Eisen, J.N., Vargas, K.J., Medeiros, A.T., Hammar, K.M., and Morgan, J.R. (2020). α-Synuclein-112 Impairs Synaptic Vesicle Recycling Consistent With Its Enhanced Membrane Binding Properties. Front. Cell Dev. Biol. 8, 405. 10.3389/fcell.2020.00405.

43. Cardo, L.F., Coto, E., de Mena, L., Ribacoba, R., Mata, I.F., Menéndez, M., Moris, G., and Alvarez, V. (2014). Alpha-synuclein transcript isoforms in three different brain regions from Parkinson’s disease and healthy subjects in relation to the SNCA rs356165/rs11931074 polymorphisms. Neurosci. Lett. 562, 45–49. 10.1016/j.neulet.2014.01.009.

44. McLean, J.R., Hallett, P.J., Cooper, O., Stanley, M., and Isacson, O. (2012). Transcript expression levels of full-length alpha-synuclein and its three alternatively spliced variants in Parkinson’s disease brain regions and in a transgenic mouse model of alpha-synuclein overexpression. Mol. Cell. Neurosci. 49, 230–239. 10.1016/j.mcn.2011.11.006.

45. Beyer, K., Domingo-Sàbat, M., Humbert, J., Carrato, C., Ferrer, I., and Ariza, A. (2008). Differential expression of alpha-synuclein, parkin, and synphilin-1 isoforms in Lewy body disease. Neurogenetics 9, 163–172. 10.1007/s10048-008-0124-6.

46. Tushev, G., Glock, C., Heumüller, M., Biever, A., Jovanovic, M., and Schuman, E.M. (2018). Alternative 3’ UTRs Modify the Localization, Regulatory Potential, Stability, and Plasticity of mRNAs in Neuronal Compartments. Neuron 98, 495–511.e6. 10.1016/j.neuron.2018.03.030.

47. Mayr, C. (2019). What are 3’ utrs doing? Cold Spring Harb. Perspect. Biol. 11. 10.1101/cshperspect.a034728.

48. Yang, X., Coulombe-Huntington, J., Kang, S., Sheynkman, G.M., Hao, T., Richardson, A., Sun, S., Yang, F., Shen, Y.A., Murray, R.R., et al. (2016). Widespread expansion of protein interaction capabilities by alternative splicing. Cell 164, 805–817. 10.1016/j.cell.2016.01.029.

49. Kelemen, O., Convertini, P., Zhang, Z., Wen, Y., Shen, M., Falaleeva, M., and Stamm, S. (2013). Function of alternative splicing. Gene 514, 1–30. 10.1016/j.gene.2012.07.083.

50. Marchese, D., Botta-Orfila, T., Cirillo, D., Rodriguez, J.A., Livi, C.M., Fernández-Santiago, R., Ezquerra, M., Martí, M.J., Bechara, E., Tartaglia, G.G., et al. (2017). Discovering the 3’ UTR-mediated regulation of alpha-synuclein. Nucleic Acids Res. 45, 12888–12903. 10.1093/nar/gkx1048.

51. Je, G., Guhathakurta, S., Yun, S.P., Ko, H.S., and Kim, Y.-S. (2018). A novel extended form of alpha-synuclein 3’UTR in the human brain. Mol. Brain 11, 29. 10.1186/s13041-018-0371-x.

52. Febbraro, F., Giorgi, M., Caldarola, S., Loreni, F., and Romero-Ramos, M. (2012). α-Synuclein expression is modulated at the translational level by iron. Neuroreport 23, 576–580. 10.1097/WNR.0b013e328354a1f0.

53. Friedlich, A.L., Tanzi, R.E., and Rogers, J.T. (2007). The 5′-untranslated region of Parkinson’s disease α-synuclein messengerRNA contains a predicted iron responsive element. Mol. Psychiatry 12, 222–223. 10.1038/sj.mp.4001937.

54. Zhou, Z.D., and Tan, E.-K. (2017). Iron regulatory protein (IRP)-iron responsive element (IRE) signaling pathway in human neurodegenerative diseases. Mol. Neurodegener. 12, 75. 10.1186/s13024-017-0218-4.

55. Merkin, J., Russell, C., Chen, P., and Burge, C.B. (2012). Evolutionary dynamics of gene and isoform regulation in Mammalian tissues. Science 338, 1593–1599. 10.1126/science.1228186.

56. Barbosa-Morais, N.L., Irimia, M., Pan, Q., Xiong, H.Y., Gueroussov, S., Lee, L.J., Slobodeniuc, V., Kutter, C., Watt, S., Colak, R., et al. (2012). The evolutionary landscape of alternative splicing in vertebrate species. Science 338, 1587–1593. 10.1126/science.1230612.

57. Bendor, J.T., Logan, T.P., and Edwards, R.H. (2013). The function of α-synuclein. Neuron 79, 1044–1066. 10.1016/j.neuron.2013.09.004.

58. Valente, A.J., Maddalena, L.A., Robb, E.L., Moradi, F., and Stuart, J.A. (2017). A simple ImageJ macro tool for analyzing mitochondrial network morphology in mammalian cell culture. Acta Histochem. 119, 315–326. 10.1016/j.acthis.2017.03.001.

59. Jiang, Y., Lee, M.-L.T., He, X., Rosner, B., and Yan, J. (2020). Wilcoxon Rank-Based Tests for Clustered Data with *R* Packageclusrank. J. Stat. Softw. 96. 10.18637/jss.v096.i06.

60. Jumper, J., Evans, R., Pritzel, A., Green, T., Figurnov, M., Ronneberger, O., Tunyasuvunakool, K., Bates, R., Žídek, A., Potapenko, A., et al. (2021). Highly accurate protein structure prediction with AlphaFold. Nature 596, 583–589. 10.1038/s41586-021-03819-2.

61. Brotzakis, Z.F., Zhang, S., and Vendruscolo, M. (2023). Alphafold prediction of structural ensembles of disordered proteins. BioRxiv. 10.1101/2023.01.19.524720.

62. Bonomi, M., Branduardi, D., Bussi, G., Camilloni, C., Provasi, D., Raiteri, P., Donadio, D., Marinelli, F., Pietrucci, F., Broglia, R.A., et al. (2009). PLUMED: A portable plugin for free-energy calculations with molecular dynamics. Comput Phys Commun 180, 1961–1972. 10.1016/j.cpc.2009.05.011.

63. Tesei, G., Schulze, T.K., Crehuet, R., and Lindorff-Larsen, K. (2021). Accurate model of liquid-liquid phase behavior of intrinsically disordered proteins from optimization of single-chain properties. Proc. Natl. Acad. Sci. USA 118. 10.1073/pnas.2111696118.

64. Mölder, F., Jablonski, K.P., Letcher, B., Hall, M.B., Tomkins-Tinch, C.H., Sochat, V., Forster, J., Lee, S., Twardziok, S.O., Kanitz, A., et al. (2021). Sustainable data analysis with Snakemake. F1000Res. 10, 33. 10.12688/f1000research.29032.2.

65. Li, H. (2018). Minimap2: pairwise alignment for nucleotide sequences. Bioinformatics 34, 3094–3100. 10.1093/bioinformatics/bty191.

66. Li, H., Handsaker, B., Wysoker, A., Fennell, T., Ruan, J., Homer, N., Marth, G., Abecasis, G., Durbin, R., and 1000 Genome Project Data Processing Subgroup (2009). The Sequence Alignment/Map format and SAMtools. Bioinformatics 25, 2078–2079. 10.1093/bioinformatics/btp352.

67. Tardaguila, M., de la Fuente, L., Marti, C., Pereira, C., Pardo-Palacios, F.J., Del Risco, H., Ferrell, M., Mellado, M., Macchietto, M., Verheggen, K., et al. (2018). SQANTI: extensive characterization of long-read transcript sequences for quality control in full-length transcriptome identification and quantification. Genome Res. 28, 396–411. 10.1101/gr.222976.117.

68. Gustavsson, E.K., Zhang, D., Reynolds, R.H., Garcia-Ruiz, S., and Ryten, M. (2022). ggtranscript: an R package for the visualization and interpretation of transcript isoforms using ggplot2. Bioinformatics 38, 3844–3846. 10.1093/bioinformatics/btac409.

69. Solntsev, S.K., Shortreed, M.R., Frey, B.L., and Smith, L.M. (2018). Enhanced Global Post-translational Modification Discovery with MetaMorpheus. J. Proteome Res. 17, 1844–1851. 10.1021/acs.jproteome.7b00873.

