## supplementary figures for "The diversity of *SNCA* transcripts in neurons, and its impact on antisense oligonucleotide therapeutics"

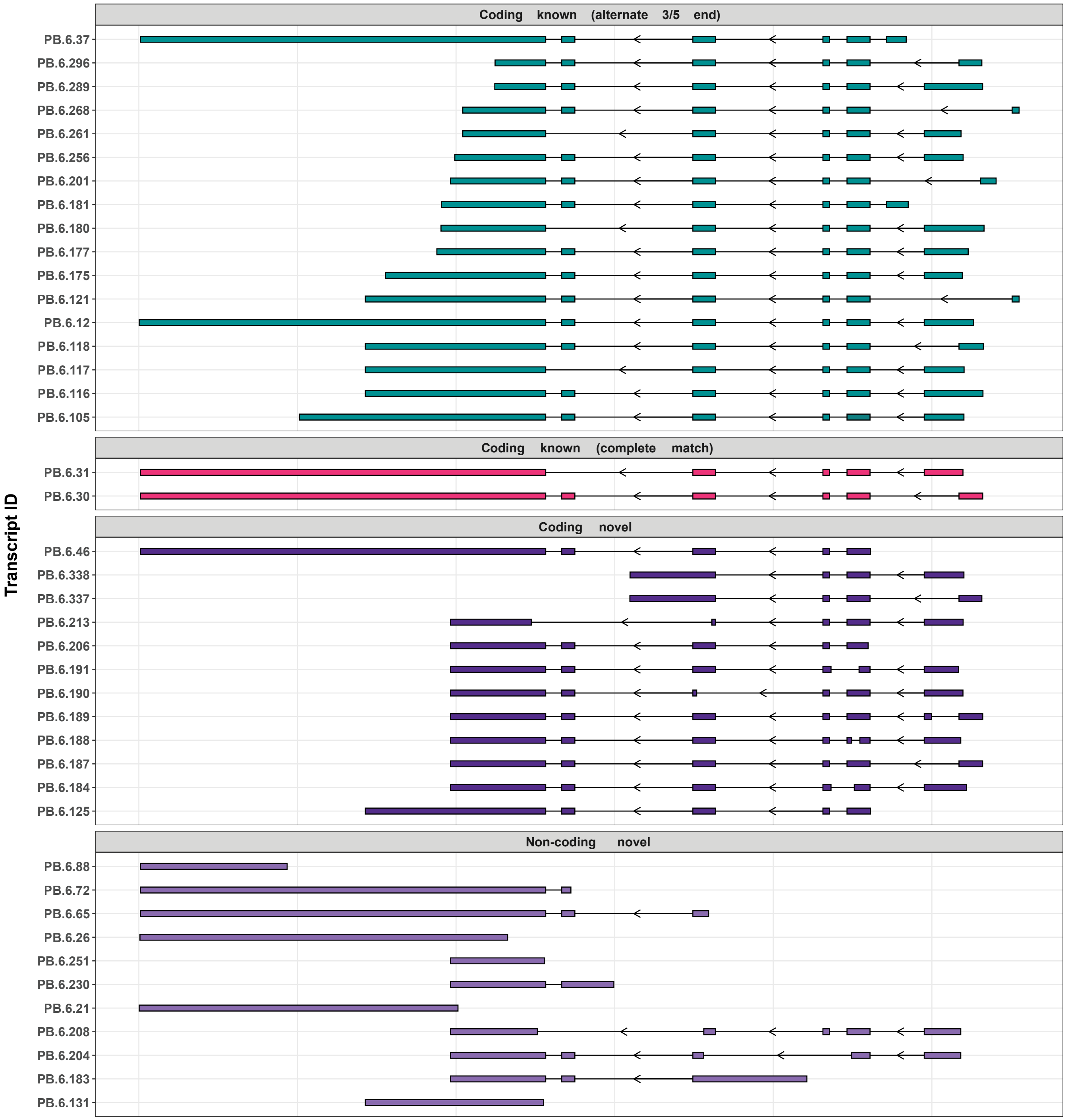


**Supplementary Fig. 2: SNCA transcript structures detected in iPSC-derived control midbrain dopaminergic neurons.** A total of identified 42 unique SNCA transcripts identified by targeted Pacific Biosciences (PacBio) long-read isoform sequencing (Iso-Seq). Coding known (alternate 3'/5' end) – if predicted to be coding & not NMD, and a full-splice match with the reference but with an alternate 3' end, 5' end or both 3' and 5' end; Coding known (complete match) – if predicted to be coding & not NMD, and a full-splice & UTR match with the reference; Coding novel – if predicted to be coding & not NMD, and not a full-splice match with the reference; and Non-coding novel – if predicted to be non-coding and not a full-splice match with the reference.

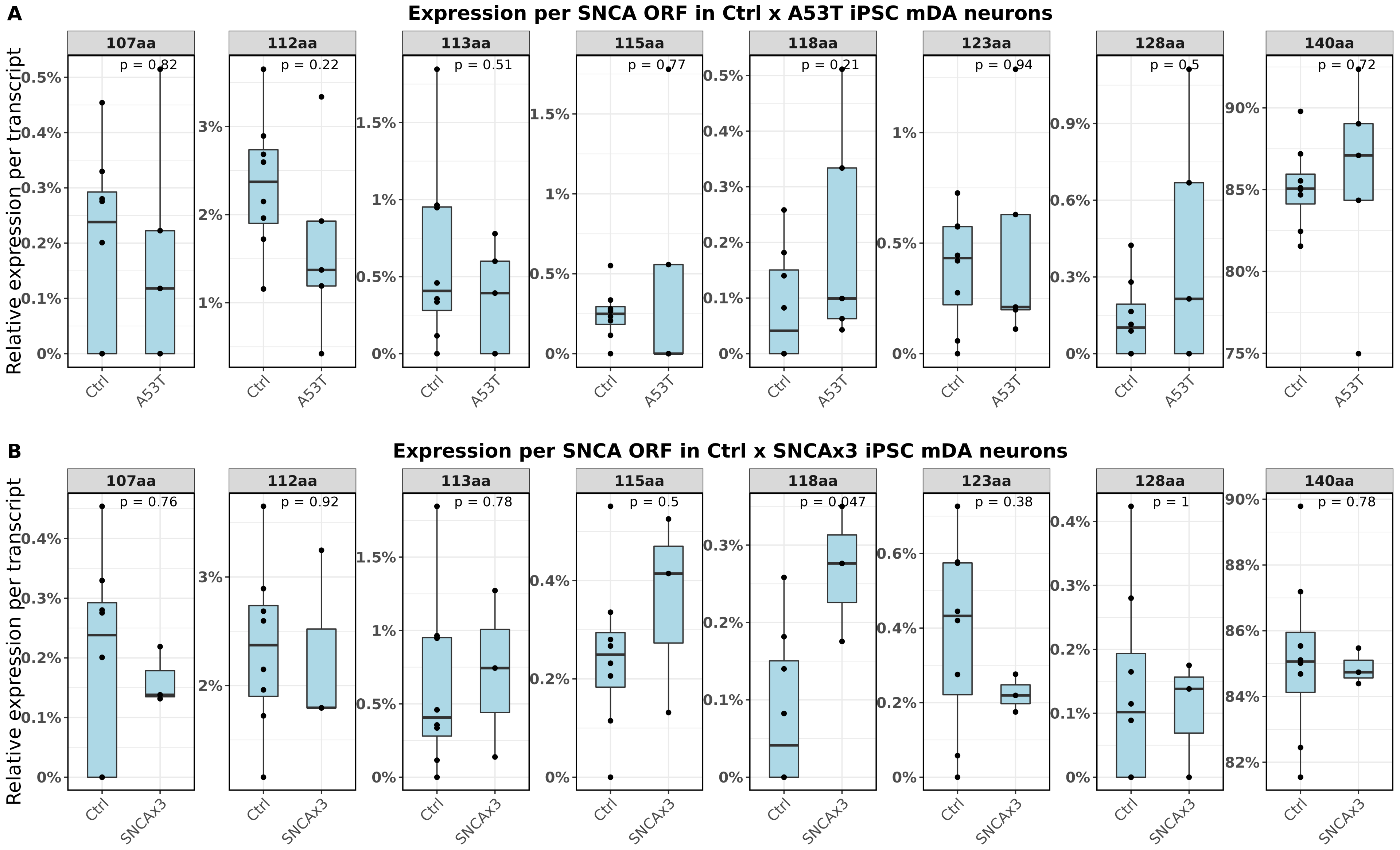


**Supplementary Fig. 4: SNCA relative transcript expression by open reading frame in control and SNCA mutant iPSC-derived mid-brain dopaminergic neurons.** (A) Relative open reading frame expression in A53T compared to control. (B) Relative open reading frame expression in SNCAx3 compared to control. A Wilcoxon test was used for statistical comparisons.

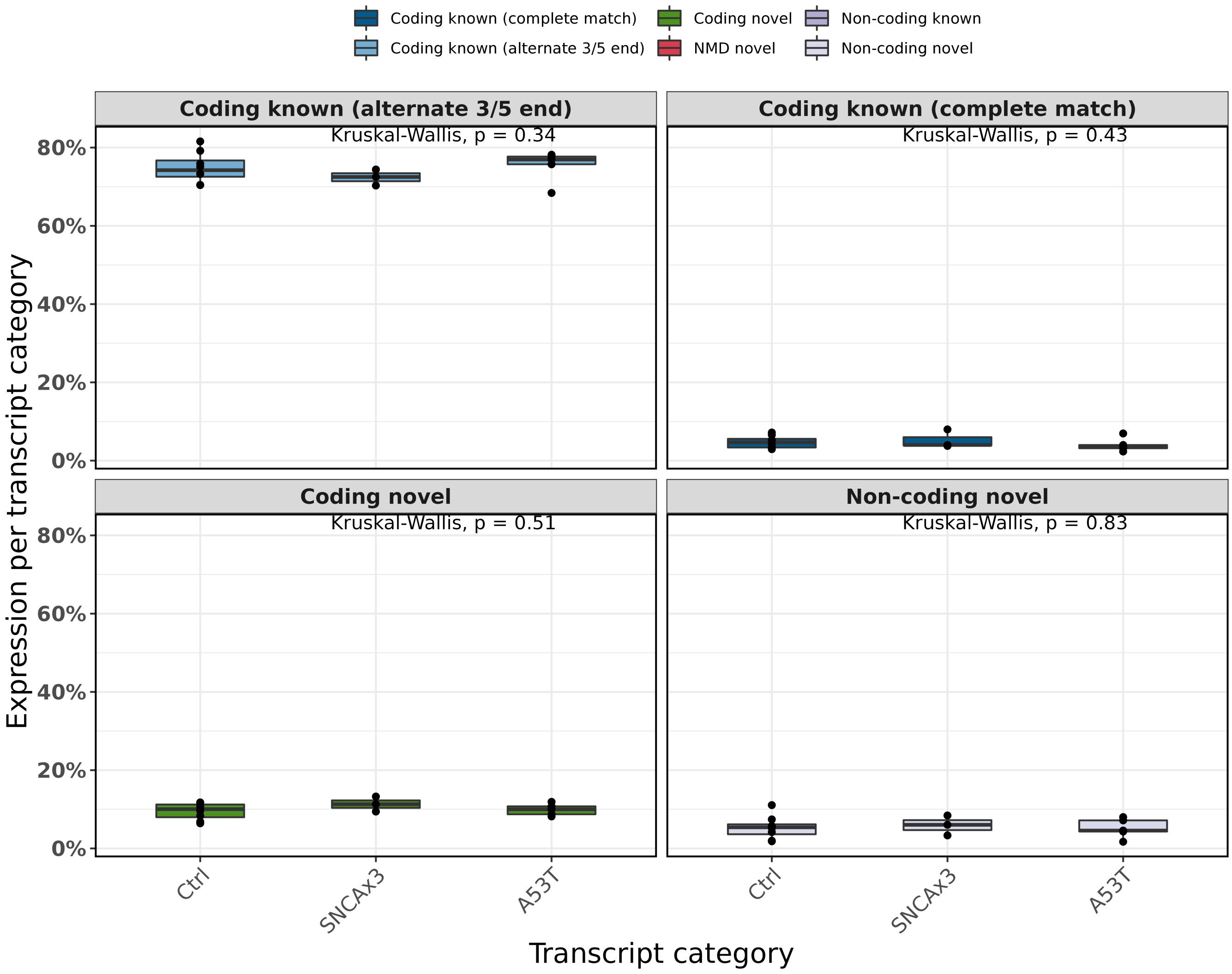


**Supplementary Fig. 3: Relative SNCA transcript expression by transcript category across genotypes.** Relative expression per transcript category: Coding known (alternate 3'/5' end) – if predicted to be coding & not NMD, and a full-splice match with the reference but with an alternate 3' end, 5' end or both 3' and 5' end; Coding known (complete match) – if predicted to be coding & not NMD, and a full-splice & UTR match with the reference; Coding novel – if predicted to be coding & not NMD, and not a full-splice match with the reference; and Non-coding novel – if predicted to be non-coding and not a full-splice match with the reference.

**A**

```
CLUSTAL O(1.2.4) multiple sequence alignment


sp|P37840|SYUA_HUMAN190          MDVFMKGLSKAKEGVVAAAEKTKQGVAEAAGKTKEGVLYVGSKTKEGVVHGVATVSKRTS 60
sp|P37840|SYUA_HUMAN213          MDVFMKGLSKAKEGVVAAAEKTKQGVAEAAGKTKEGVLYVGSKTKEGVVHGVATVAEKTK 60
sp|P37840|SYUA_HUMAN338          MDVFMKGLSKAKEGVVAAAEKTKQGVAEAAGKTKEGVLYVGSKTKEGVVHGVATVAEKTK 60
sp|P37840|SYUA_HUMAN191          MDVFMKGLSKAKEG----------------------VVFVGSKTKEGVVHGVATVAEKTK 38
sp|P37840|SYUA_HUMAN184          MDVFMKGLSKAKEGVVAAAEKTKQG------------VFVGSKTKEGVVHGVATVAEKTK 48
sp|P37840|SYUA_HUMANCanonical    MDVFMKGLSKAKEGVVAAAEKTKQGVAEAAGKTKEGVLYVGSKTKEGVVHGVATVAEKTK 60
sp|P37840|SYUA_HUMAN188          MDVFMKGLSKAKE-----------------GKTKEGVLYVGSKTKEGVVHGVATVAEKTK 43
                                 *************                     ::****************:::*.

sp|P37840|SYUA_HUMAN190          WARMKKEPHRKEFW-KICLWILTMRLMKCL-LRKGIKTTNLKPKKYLC---SQFLEIC-- 113
sp|P37840|SYUA_HUMAN213          EQVLSSNVPSHDI----SQS--FYSVSRSLPSAV--I------EVSVPAPTQHFGASLSL 106
sp|P37840|SYUA_HUMAN338          EQVTNVGGAVVTGVTAVAQK--TVEGAGSIAAATGFVKKDQLGKVWLCTFCVTFISW--- 115
sp|P37840|SYUA_HUMAN191          EQVTNVGGAVVTGVTAVAQK--TVEGAGSIAAATGFVKKDQLGKNEEGAPQEGILEDMPV 96
sp|P37840|SYUA_HUMAN184          EQVTNVGGAVVTGVTAVAQK--TVEGAGSIAAATGFVKKDQLGKNEEGAPQEGILEDMPV 106
sp|P37840|SYUA_HUMANCanonical    EQVTNVGGAVVTGVTAVAQK--TVEGAGSIAAATGFVKKDQLGKNEEGAPQEGILEDMPV 118
sp|P37840|SYUA_HUMAN188          EQVTNVGGAVVTGVTAVAQK--TVEGAGSIAAATGFVKKDQLGKNEEGAPQEGILEDMPV 101
                                     .            .          .:              :         :

sp|P37840|SYUA_HUMAN190          ----------------------      113
sp|P37840|SYUA_HUMAN213          K---------------------      107
sp|P37840|SYUA_HUMAN338          ----------------------      115
sp|P37840|SYUA_HUMAN191          DPDNEAYEMPSEEGYQDYEPEA      118
sp|P37840|SYUA_HUMAN184          DPDNEAYEMPSEEGYQDYEPEA      128
sp|P37840|SYUA_HUMANCanonical    DPDNEAYEMPSEEGYQDYEPEA      140
sp|P37840|SYUA_HUMAN188          DPDNEAYEMPSEEGYQDYEPEA      123
```

**B**

| Isoform | Unique tryptic sequences | Detection summary |
|---|---|---|
| 190 | ICLWILTMR, YLCSQFLEIC* | YLCSQFLEIC* detected in tissue IP and tissue lysate |
| 213 | EQVLSSNVPSHDISQSFYSVSR, SLPSAVIEVSVPAPTQHFGASLSLK* | Not detected |
| 338 | VWLCTFCVTFISWA* | Not detected |
| 184 | QGVFVGSKTK | Not detected |
| 191 | EGVVFVGSK | Detected in tissue IP and but not confidently in tissue lysate |
| 188 | No identifiable unique sequences | |

* C-terminal sequence

**Supplementary Fig. 5:** (A) Multiple sequence alignment of predicted isoform sequences with canonical αSyn sequence. Isoform unique tryptic sequences are boxed in red and a common αSyn sequence used to detect wild-type and majority of isoforms is boxed in blue. (B) List of identified tryptic unique peptides used to detect novel α Syn isoforms and summary of detection in tissue immunopreceipitation as well as whole tissue lysate

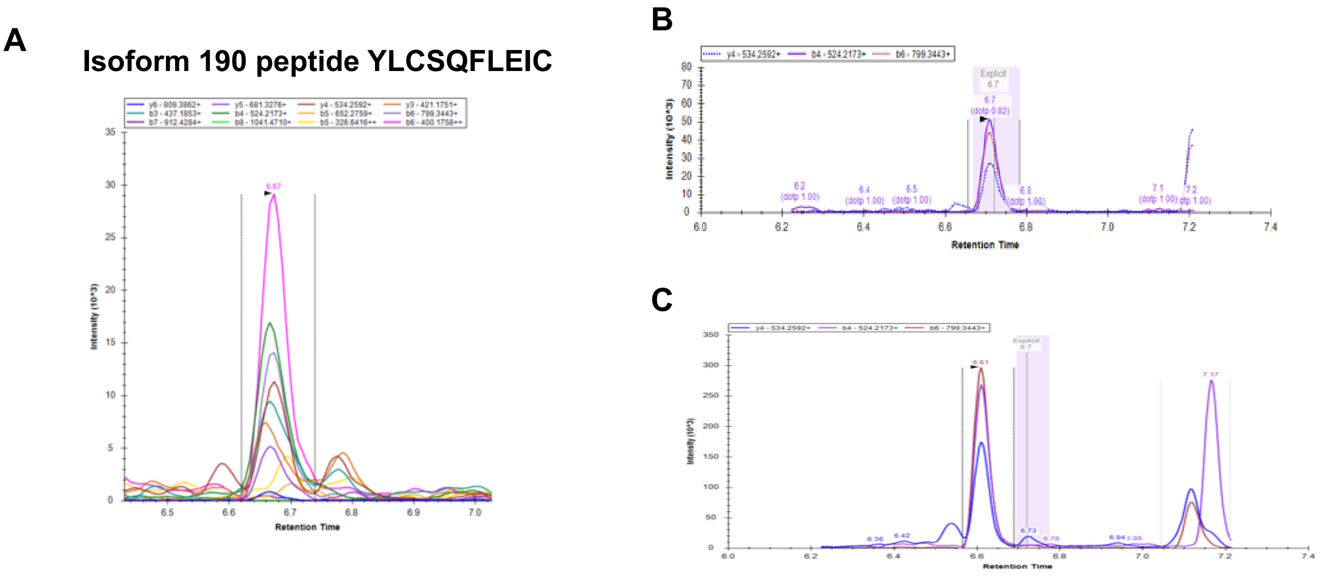


**Supplementary Fig. 6: Detection of isoform 190 using Multiple Reaction Monitoring analysis.** Overlaid chromatogram shows multiple transitions for 190 peptide YLCSQFLEIC (A) spiked into brain lysate (B) endogenous peptide detected in tissue IP and (C) in brain tissue lysate.

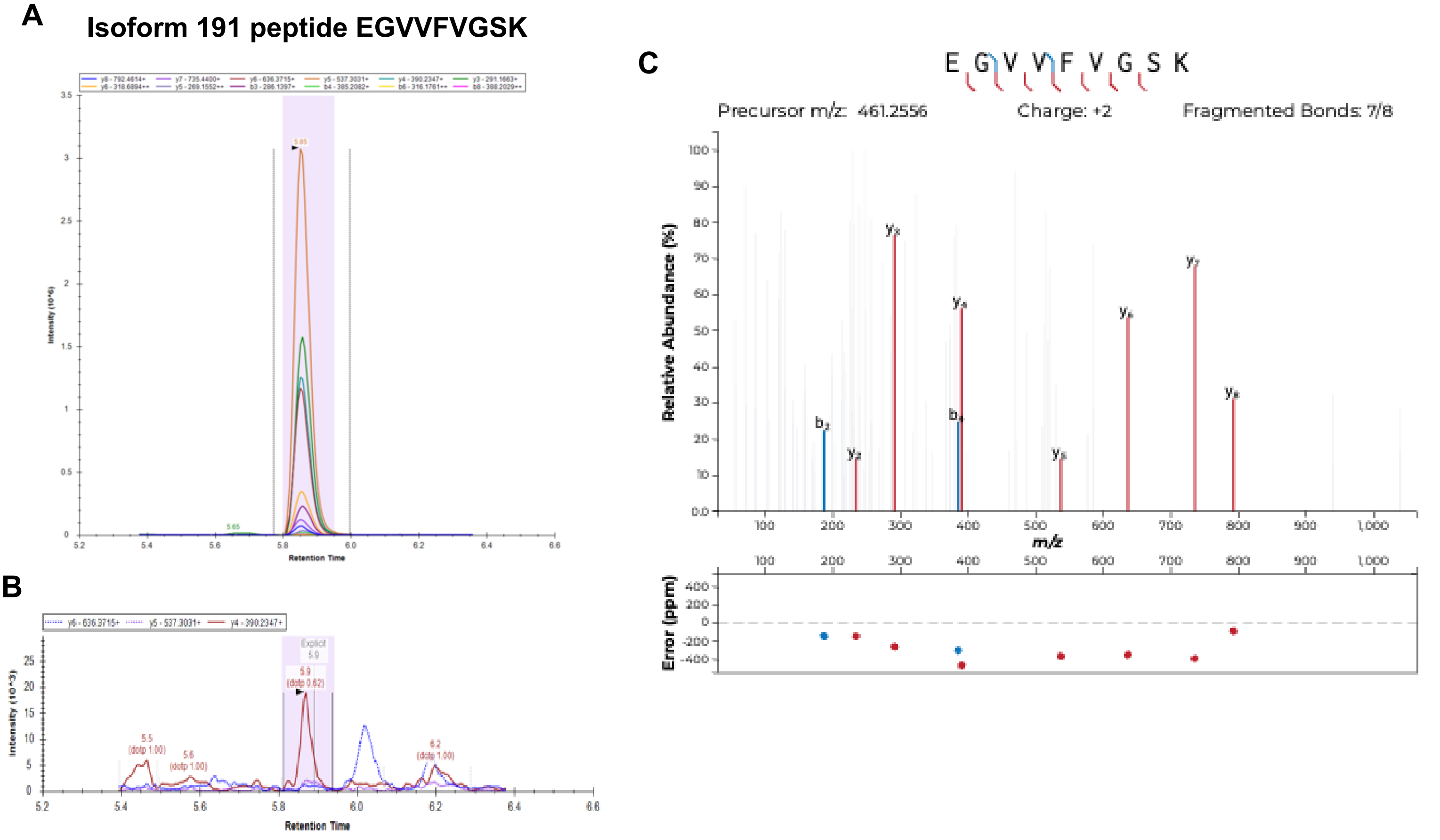


**Supplementary Fig. 7: Detection of isoform 191 using Multiple Reaction Monitoring analysis.** Overlaid chromatogram shows multiple transitions for 191 peptide EGVVFVGSK (A) spiked into brain lysate (B) endogenous peptide detected in tissue IP and (C) shows EGVVFVGSK fragment ion spectra with mass error

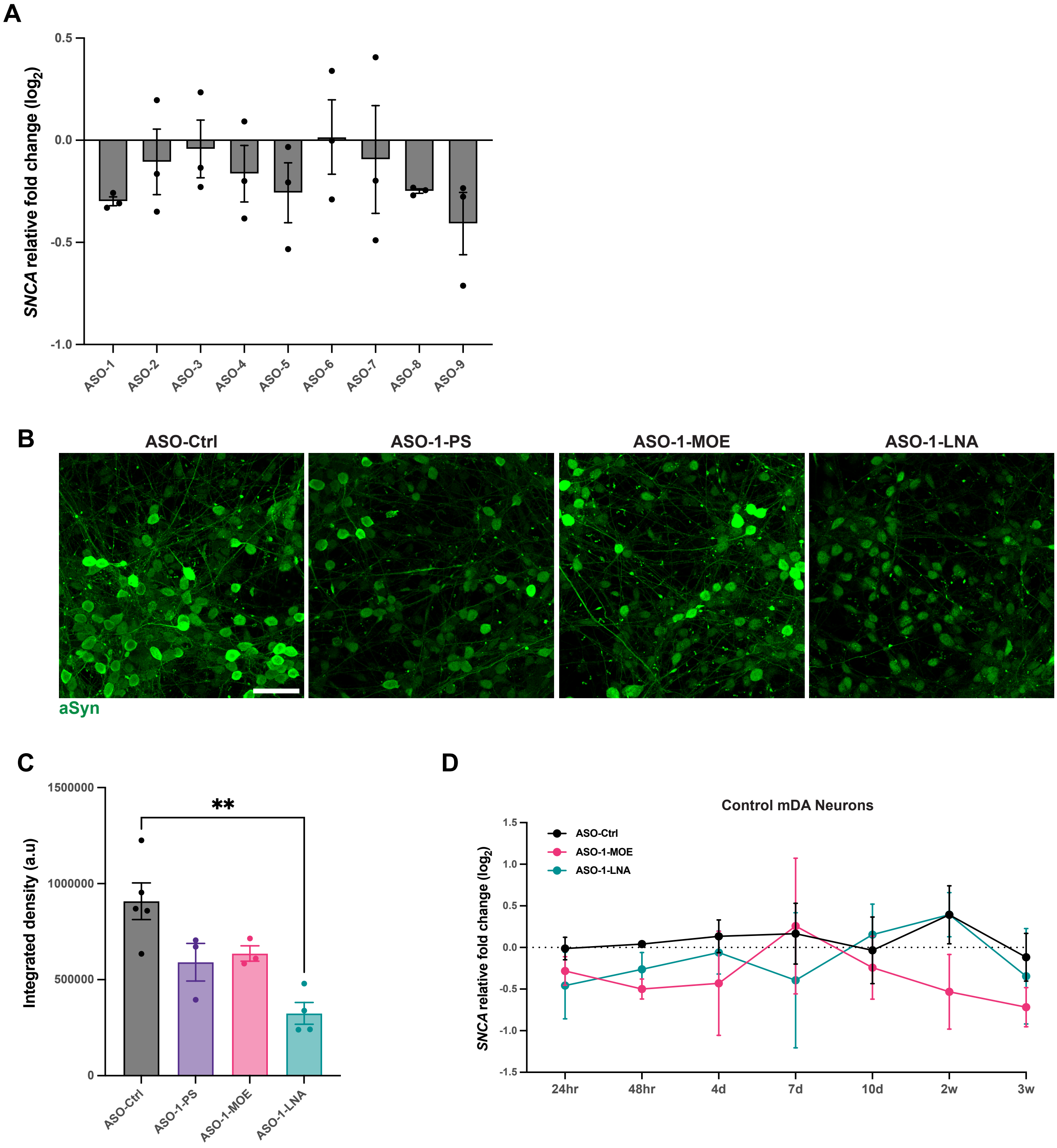


**Supplementary Fig. 8: ASO design and optimisation.** (A) Quantitative PCR showing mRNA expression SH-SY5Ys treated with ASOs for 48 hours relative to a non-targeting control ASO. (B) Representative ICC total αSyn in mDA neurons treated with ASO-Ctrl, ASO-1-PS, ASO-1-MOE, ASO-1-LNA for 24 hours. (G) ICC quantification of the integrated density of αSyn (**P < 0.01, one-way ANOVA with Tukey's multiple comparison test). (D) Quantitative PCR data showing *SNCA* expression at multiple timepoints after transfection in control iPSC-derived mDA neurons (n = 3 donor lines).

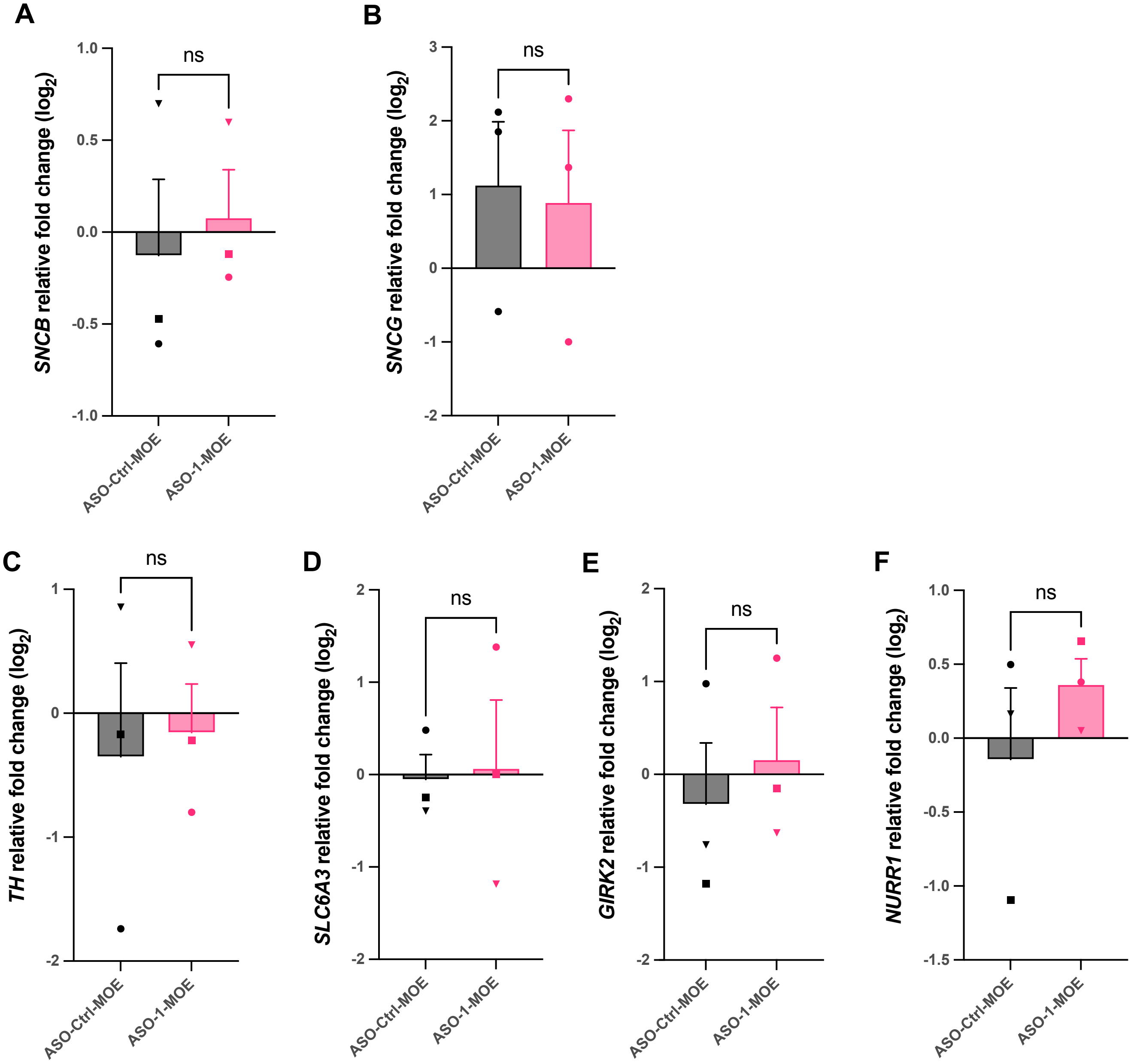


**Supplementary Fig. 9: ASO specificity and impact impact on neuronal differentiation.** (A) Quantitative PCR showing mRNA expression of (A) *SNCB* and (C) *SNCG* in mDA neurons 10 days after treatment with ASO-Ctrl-MOE and ASO-1-MOE. Quantitative PCR showing mRNA expression of the midbrain markers (C) *TH*, (D) *SLC6A3*, (E) *GIRK2*, and (F) *NURR1* in mDA neurons 10 days after treatment with ASO-1-MOE and ASO-1-MOE

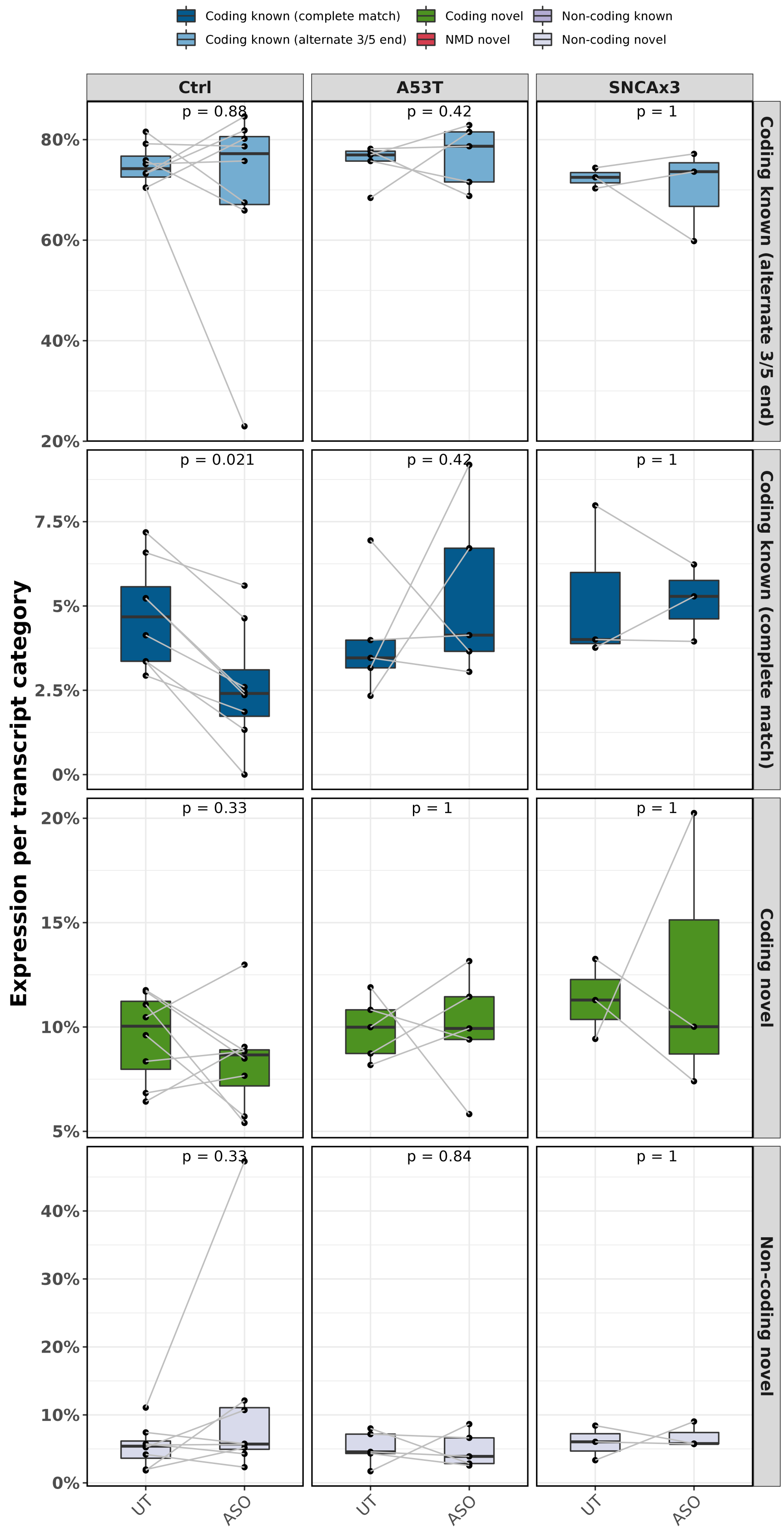


**Supplementary Fig. 10: Relative SNCA transcript expression by transcript category in ASO treated mDA neurons across genotypes.** Relative expression per transcript category in samples untreated (UT) and treated with antisense oligonucleotide targeting the 3'UTR of SNCA (ASO): Coding known (alternate 3'/5' end) – if predicted to be coding & not NMD, and a full-splice match with the reference but with an alternate 3' end, 5' end or both 3' and 5' end; Coding known (complete match) – if predicted to be coding & not NMD, and a full-splice & UTR match with the reference; Coding novel – if predicted to be coding & not NMD, and not a full-splice match with the reference; and Non-coding novel – if predicted to be non-coding and not a full-splice match with the reference. A Wilcoxon test was used for statistical comparisons and connecting lines represent samples with and without treatment.

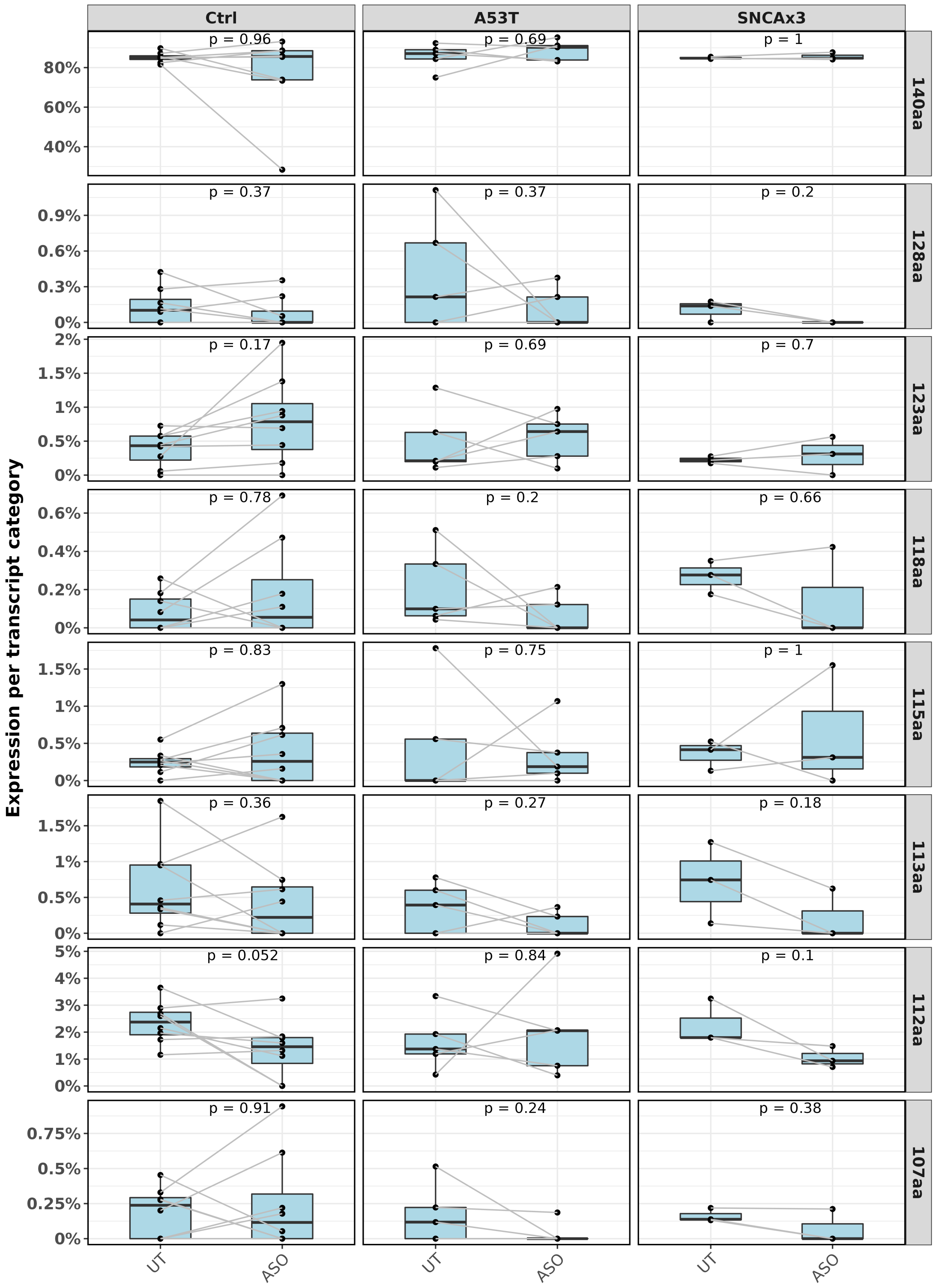


**Supplementary Fig. 11: SNCA relative transcript expression by open reading frame in control and SNCA mutant iPSC-derived midbrain dopaminergic neurons.** Relative expression per open reading frame in samples untreated (UT) and treated with antisense oligonucleotide targeting the 3'UTR of SNCA (ASO) in Ctrl, A53T and SNCAx3 dopaminergic neurons. A Wilcoxon test was used for statistical comparisons and connecting lines represent samples with and without treatment.

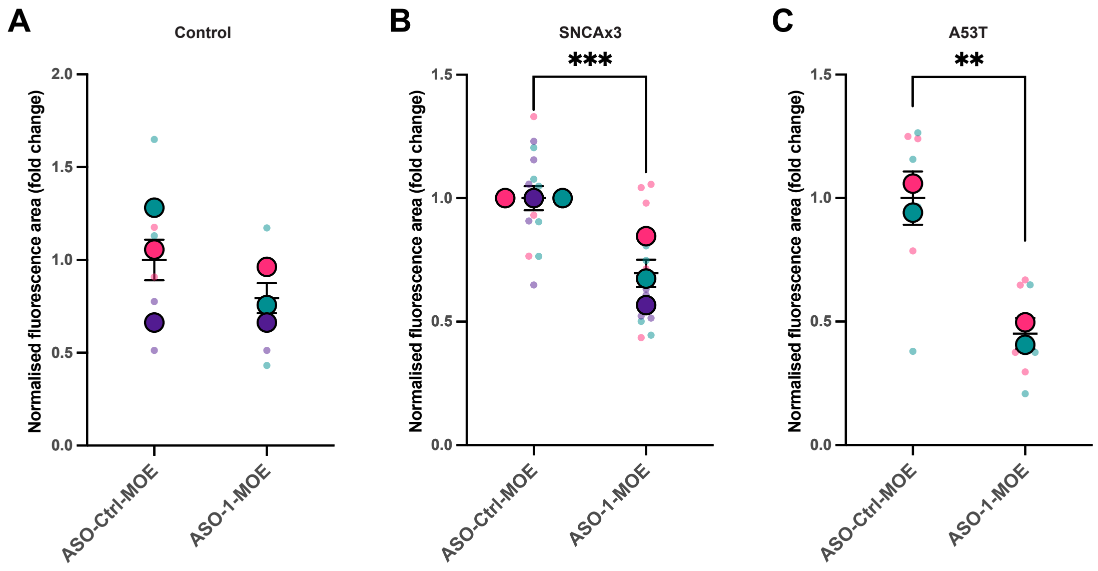


**Supplementary Fig. 12: *SNCA*-targeting ASO reduces αSyn protein 10 days after transfection in mDA neurons.** (A) Quantification of the fluorescence area of total αSyn in (A) control, (B) SNCAx3, and (C) A53T mDA neurons with ASO treatment (control = 3 donor lines, SNCAx3 = 3 independent experiments, A53T = 2 donor lines; $^{**}P < 0.01$, $^{***}P < 0.001$, clustered Wilcoxon rank sum test).

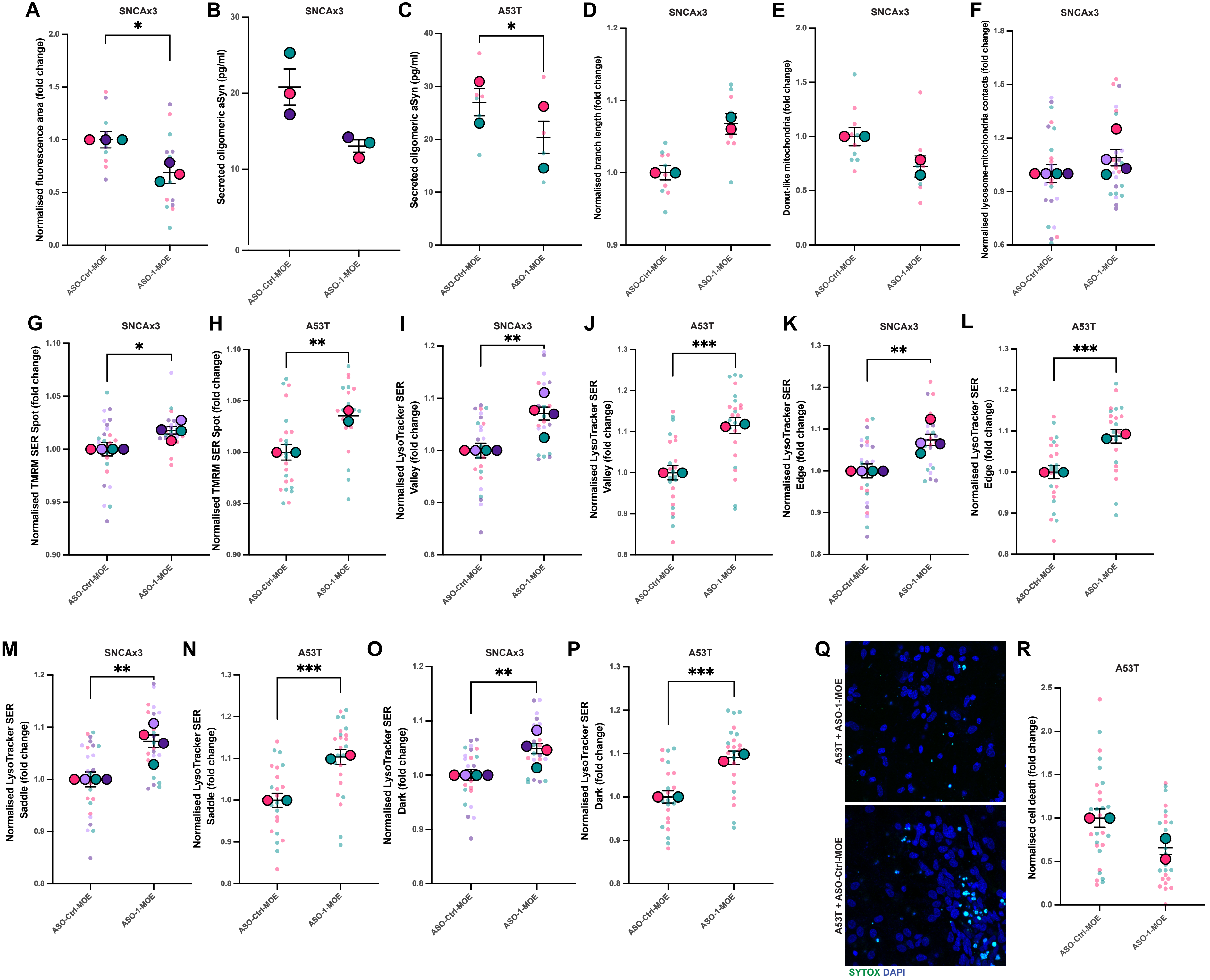
A
SNCAx3
Normalised fluorescence area (fold change)
ASO-Ctrl-MOE
ASO-1-MOE
B
SNCAx3
Secreted oligomeric aSyn (pg/ml)
C
A53T
Secreted oligomeric aSyn (pg/ml)
D
SNCAx3
Normalised branch length (fold change)
E
SNCAx3
Donut-like mitochondria (fold change)
F
SNCAx3
Normalised lysosome-mitochondria contacts (fold change)
G
SNCAx3
Normalised TMRM SER Spot (fold change)
H
A53T
Normalised TMRM SER Spot (fold change)
I
SNCAx3
Normalised LysoTracker SER Valley (fold change)
J
A53T
Normalised LysoTracker SER Valley (fold change)
K
SNCAx3
Normalised LysoTracker SER Edge (fold change)
L
A53T
Normalised LysoTracker SER Edge (fold change)
M
SNCAx3
Normalised LysoTracker SER Saddle (fold change)
N
A53T
Normalised LysoTracker SER Saddle (fold change)
O
SNCAx3
Normalised LysoTracker SER Dark (fold change)
P
A53T
Normalised LysoTracker SER Dark (fold change)
Q
A53T + ASO-1-MOE
A53T + ASO-Ctrl-MOE
SYTOX DAPI
R
A53T
Normalised cell death (fold change)

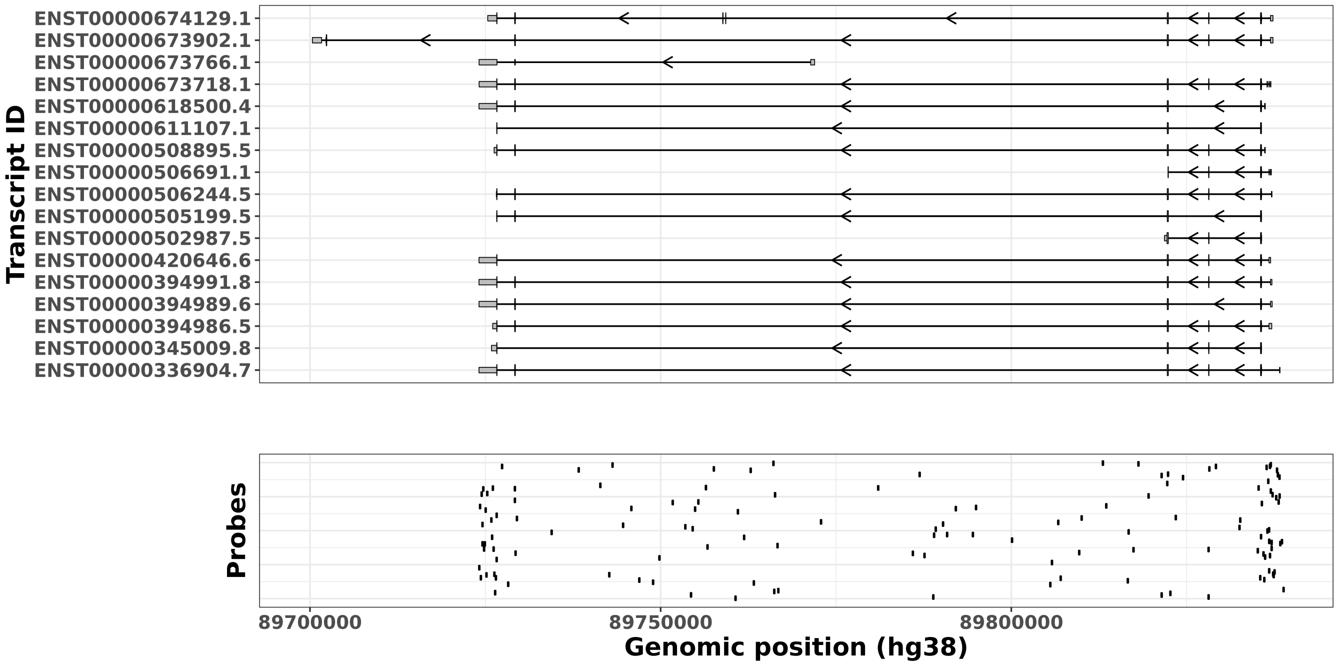


**Supplementary Fig 1: SNCA hybridization probe design.** Enrichment of SNCA cDNAs were done using 111 biotinylated IDT lockdown hybridization probes, designed against exonic (n = 41) and intronic (n = 70) genic regions of SNCA according to annotation (GENCODEv38). The top panel show annotated transcripts according to GENCODEv38 and the bottom panel shows the location of the hybridization on chr4 (hg38).

**Supplementary Fig. 13: ASO-1-MOE reduces PD-associated cellular phenotypes.** (A) Quantification of the fluorescence area of aggregated αSyn in ASO treated SNCAx3 mDA neurons (n = 3 independent differentiations; *$P$< 0.05, paired t-test). (B) Quantification of secreted oligomeric αSyn by ELISA in SNCAx3 mDA neurons (n = 3 independent differentiations; *$P$< 0.05, paired t-test). (C) Quantification of secreted oligomeric αSyn by ELISA in A53T mDA neurons with ASO treatment (n = 2 donor lines each from 3 independent differentiations; *$P$< 0.05, clustered Wilcoxon rank sum test). (D) Quantification of the mitochondrial branch length in SNCAx3 mDA neurons with ASO treatment using MINA (paired t-test). (E) Quantification of donut-like mitochondria using MINA (paired t-test). (F) Quantification of the number of mitochondria-lysosome contacts in SNCAx3 mDA neurons with ASO treatment (clustered Wilcoxon rank sum test). (G-H) Quantification of TMRM SER spot in SNCAx3 and A53T mDA neurons with ASO treatment (SNCAx3 = 4 independent differentiations, A53T = 2 donor lines across 4 independent differentiations; *$P$< 0.05, **$P$< 0.01, clustered Wilcoxon rank sum test). Quantification of the lysosomal textural features (I-J) SER Valley, (K-L) SER Edge, (M-N) SER Saddle, and (O-P) SER Dark (SNCAx3 = 4 independent differentiations, A53T = 2 donor lines across 4 independent differentiations; *$P$< 0.05, **$P$< 0.01, ***$P$< 0.001, clustered Wilcoxon rank sum test). (Q) Live-cell images depicting dead cells in SNCAx3 mDA neurons with and without ASO treatment using the dye SYTOX green. (R) Quantification of the proportion of dead cells in A53T mDA neurons with and without ASO treatment (paired t-test).
